# SARS-CoV-2 RBD *in vitro* evolution follows contagious mutation spread, yet generates an able infection inhibitor

**DOI:** 10.1101/2021.01.06.425392

**Authors:** Jiří Zahradník, Shir Marciano, Maya Shemesh, Eyal Zoler, Jeanne Chiaravalli, Björn Meyer, Yinon Rudich, Orly Dym, Nadav Elad, Gideon Schreiber

## Abstract

SARS-CoV-2 is continually evolving, with more contagious mutations spreading rapidly. Using *in vitro* evolution to affinity maturate the receptor-binding domain (RBD) of the spike protein towards ACE2 resulted in the more contagious mutations, S477N, E484K, and N501Y, to be among the first selected, explaining the convergent evolution of the “European” (20E-EU1), “British” (501.V1),”South African” (501.V2), and ‘‘Brazilian” variants (501.V3). Plotting the binding affinity to ACE2 of all RBD mutations against their incidence in the population shows a strong correlation between the two. Further *in vitro* evolution enhancing binding by 600-fold provides guidelines towards potentially new evolving mutations with even higher infectivity. For example, Q498R epistatic to N501Y. Nevertheless, the high-affinity RBD is also an efficient drug, inhibiting SARS-CoV-2 infection. The 2.9Å Cryo-EM structure of the high-affinity complex, including all rapidly spreading mutations, provides a structural basis for future drug and vaccine development and for *in silico* evaluation of known antibodies.

## Introduction

SARS-CoV-2, which causes COVID-19, resulted in an epidemic of global reach. It infects people through inhalation of viral particles, airborne, in droplets, or by touching contaminated surfaces. Structural and functional studies have shown that a single receptor-binding domain (RBD) of the SARS-CoV-2 homotrimer spike glycoprotein interacts with ACE2, which serves as its receptor^1,2^. Its binding and subsequent cleavage by the host protease TMPRSS2 results in the fusion between the cell and viral membranes and cell entry^1^. Blocking the ACE2 receptors by specific antibodies voids viral entry^1,3,4^ *In vitro* binding measurements have shown that SARS-CoV-2 S-protein binds ACE2 with ~10 nM affinity, which is about 10-fold tighter than the binding of the SARS-CoV S-protein^2,3,5^. It was suggested that this is, at least partially, responsible for its higher infectivity^6^. Recently evolved SARS-CoV-2 mutations in the spike protein’s RBD have further strengthened this hypothesis. The “British” mutation (N501Y) was suggested from deep sequencing mutation analysis to enhance binding to ACE2^6^. The “South African” mutation, which includes three altered residues in the ACE2 binding site (K417N, E484K, and N501Y) spread extremely rapidly, becoming within weeks the dominant lineage in the Eastern Cape and Western Cape Provinces^7^. The “Brazilian” variant (P.1), independently fixing K417T, E484K, and N501Y mutations similar to the “South African” variant, is spreading rapidly from the Amazon region^8^. Another variant enhancing SARS-CoV-2 infectivity is S477N, which became dominant in many regions^9^. Recently, several efficient vaccines, based on presenting the spike protein or by administrating an inactivated virus were approved for clinical use^10^. Still, due to less than 100% protection, particularly for high-risk populations and the continuously mutating virus, drug development should continue. Potential therapeutic targets blocking viral entry include molecules that block the spike protein, the TMPRSS2 protease, or the ACE2 receptor^11^. Most prominently, multiple high-affinity neutralizing antibodies have been developed^12^. Alternatives to the antibodies, the soluble forms of the ACE2 protein^13^ or engineered parts or mimics have also been shown to work^14,15^. TMPRSS2 inhibitors were already previously developed, and are repurposed for anti-COVID-19 treatment^1^. The development of molecules blocking the ACE2 protein did not receive as much attention as the other targets. One potential civet with this approach is the importance of the ACE2 activities, both as a carboxypeptidase, removing a single C-terminal amino acid from Ang II to generate Ang-(1-7) and regulating amino acid transport and pancreatic insulin secretion^16,17^ These functions, which could be hampered by an inhibitor, are important in regulating blood pressure and inflammation, which downregulation relates to increased COVID-19 severity. Dalbavancin is one drug that blocks the spike protein–ACE2 interaction, however with low affinity (~130 nM)^18^.

Notably, the RBD domain itself can be used as a competitive inhibitor of the ACE2 receptor binding site. However, for this to work, its affinity has to be significantly optimized, to reach pM affinity. We have recently developed an enhanced strategy for yeast display, based on C and N-terminal fusions of extremely bright fluorescent moieties that can monitor expression at minute levels, allowing for selection to proceed down to pM bait concentrations^19^. The affinity maturated RBDs were strikingly similar to highly contagious virus variants, which motivated us to study the observed phenomenon in greater detail.

### RBD domain affinity maturation recapitulates multiple steps in the virus evolution

For *in vitro* evolution, we took advantage of an enhanced yeast surface display protocol recently developed by us, using two different detection strategies, eUnaG2 and DnbALFA^19^. Preceding library construction, we tested varied length of the RBD for optimal surface expression and stability (Table S1). Subsequently, we chose RBDcon2 for yeast display and RBDcon3 for protein expression (Supplementary Material Text). Libraries were constructed in a step-wise manner: first the S (stability-enhancing), followed by B (ACE2 binding, B3-B5), and finally B6(FA) (fast association) (Fig. 1). All libraries were constructed by error-prone PCR random mutagenesis of the RBD, introducing 1-5 mutations per clone (Fig. 1). Libraries S1 and S2 converged towards the I358F mutation, with the phenylalanine residue fitting into the hydrophobic pocket formed in the RBD domain (Fig. 1 and S1). This mutation nearly doubled the fluorescence signal intensity and was used to construct library B3 (Fig. 1), where mutations were limited to residues G431-K528 (Fig. 1). The B3 library was expressed at 37 °C to keep the pressure on protein stability, and selected by FACS sort against decreasing concentration of ACE2 labeled with CF^®^640R succinimidyl ester (1000, 800, and 600 pM; 4 h of incubation). Library enrichment was achieved by selecting the top 3% of binding cells in the first round and in subsequent rounds, the top 0.1 – 1% yeast cells (Fig. 1 and Fig. S2). Plasmid DNA was isolated from selected yeasts of the sorted library and used for *E. coli* cell transformation and the preparation of a new library (B4). This approach, which mimics natural virus evolution, enriches the subsequent library with multiple selected mutations and purges non-beneficial mutations, enabling the screening of wider sequence-space and epistatic mutations, with multiple trajectories being sampled. 30 single colony isolates were used for sequencing to monitor the enrichment process and subsequently for binding affinity screening (Fig. S3). Analysis of the selected B3 library yielded two dominant mutations appearing at > 70 % of clones: E484K and N501Y. In addition, multiple minor mutations: V483E, N481Y, I468T, S477N, N448S, and F490S were found (Table 1 and S2). The analysis of library B4 (selected against 600, 400, and 200 pM ACE2) showed the absolute domination of E484K and N501Y. Besides the dominant clones, mutations N460K, Q498R, and S477N rose to frequencies > 20 %, and new minor populated mutations were identified: G446R, I468V, T478S, F490Y, and S494P. We choose clones with different mutation profiles to validate our results, expressed them in Expi293F™ cells, and subjected them for further analyses (Figs. 2B and D, S4, and Tables 1 and 2).

**Fig. 1.**
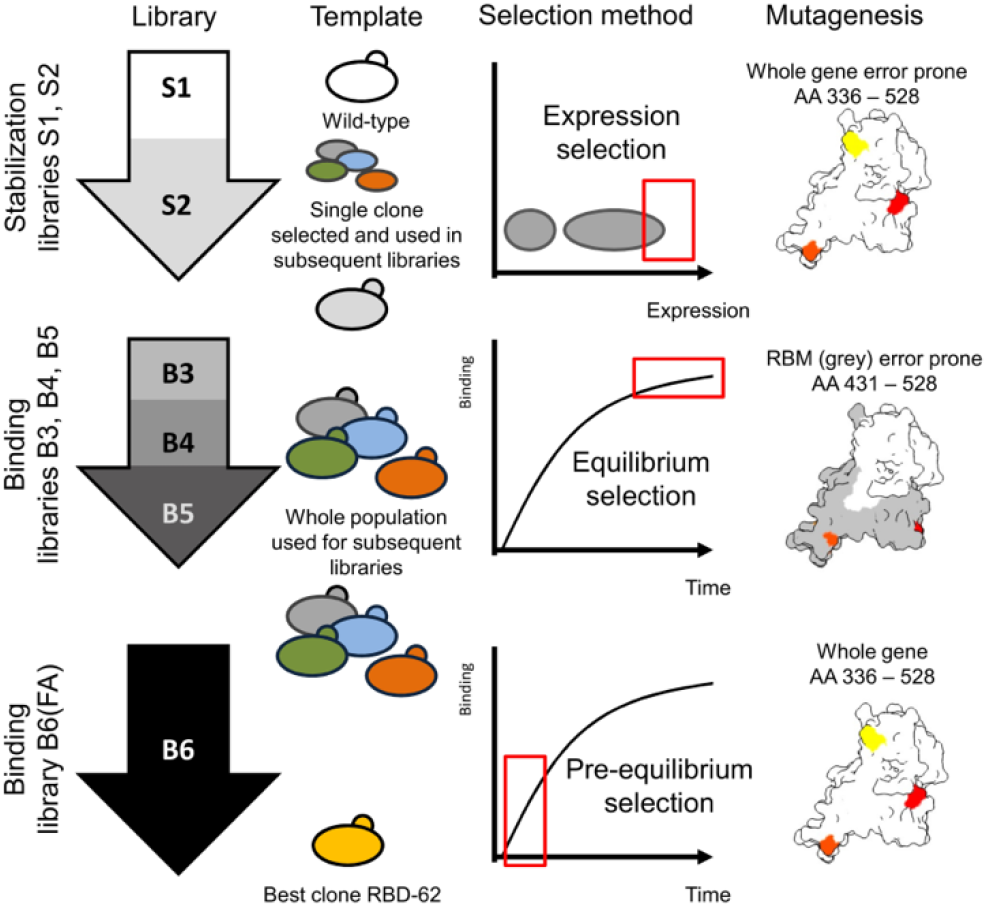
SARS-CoV2 RBD *in vitro* selection. Six consecutive libraries were created by gene error-prone PCR (1 – 5 mutations per gene) to increase RBD domain stability and binding to the ACE2 receptor. Libraries S1 and S2 were selected for higher expression at 30 °C 37 °C respectively (see Fig. S2). A single clone, I358F was identified (Fig. S1). Based on this stabilized RBD, we created library B3 to B5, randomizing residues 431-528. From each selected library the pool of enriched clones was used as a template for the subsequent library. The last library B6(FA) was created by pre-equilibrium selection of randomly mutated residues 336-528. All libraries were sequenced and monitored for affinity and stability (Table S2).

**Fig. 2.**
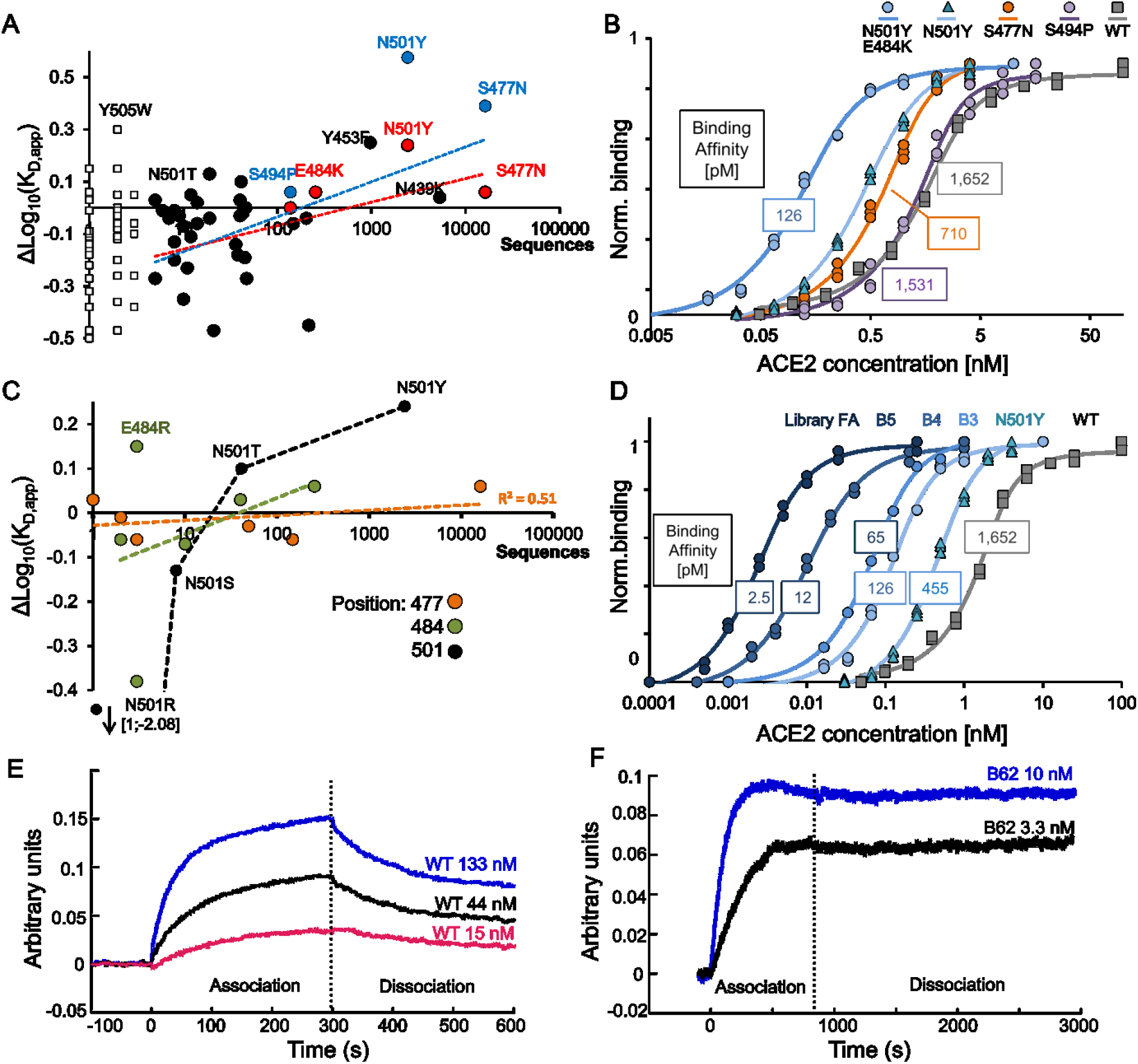
*In vitro* evolution of spike protein RBD using yeast display and the emergence of mutations in SARS-CoV-2 over time. (A) The relation between apparent binding affinity against ACE2 of single amino-acid mutations as calculated from deep-mutational scanning of the RBD domain^6^ is plotted against their frequency in the GISAID database^22^. Red is for prevalent mutations, black for others, and empty squares for mutations with <5 sequences. Blue dots are values calculated from our experimental binding titration curves shown in (B). The differences between red and blue dots of the same mutation relate to the difference between deep-mutational scanning estimates and experimental titration curves. (C) Affinity changes calculated from deep-mutational scanning plotted against frequency in the population as in (A), but for different mutations at given positions (477, 484, and 501). (D) Binding titration curves for the best binding variant in each yeast display affinity maturation library and N501Y mutation for comparison. The best clone from the first library (B3) is the E484K/N501Y double mutant, found in the “South African” and ““Brazilian”“ variants. The binding parameters of additional clones from each library is shown in Fig. S3 and Table S2. (F, G) Octet RED96 System binding sensorgrams for RBD-WT (E) and RBD-62 (F).

**Table 1 –.** RBD domain mutations selected by *in vitro* evolution for higher affinity versus those spreading in SARS-CoV-2.

| ***In vitro* evolution** |  |  |  |  |  |  |  |  |  |  |  |  |  |  |  |  |  |  |
| --- | --- | --- | --- | --- | --- | --- | --- | --- | --- | --- | --- | --- | --- | --- | --- | --- | --- | --- |
| RBD residue* | 354 | 358 | 445 | 446 | 448 | 460 | 468 | 470 | 477 | 478 | 481 | 483 | 484 | 490 | 493 | 494 | 498 | 501 |
| RBD wild-type | N | I | V | G | N | N | I | T | S | T | N | V | E | F | Q | S | Q | N |
| Library S2 |  | F |  |  |  |  |  |  |  |  |  |  |  |  |  |  |  |  |
| Library B3 |  | F |  |  | S |  | T |  | N |  | Y | E | K | S |  |  |  | Y |
| Library B4 |  | F |  | R |  | K | V |  | N | S |  |  | K | Y |  | P | R | Y |
| Library B5 |  | F |  | R |  | K | V |  | N |  |  |  | K | Y | H | P | R | Y |
| Library B6(FA) | E | F | K |  |  | K | T | M | N |  |  |  | K |  |  | P | R | Y |
| Clone B62 |  | F | K |  |  | K | T | M | N |  |  |  | K |  |  |  | R | Y |
| SARS-CoV2 var. | parental/lineage** |  |  |  |  | 460 | 468 | 470 | 477 | 478 | 481 | 483 | 484 | 490 | 493 | 494 | 498 | 501 |
| Europe | 20E/EU1 |  |  |  |  |  |  |  | N |  |  |  |  |  |  |  |  |  |
| British | 20I/501Y.V1 |  |  |  |  |  |  |  |  |  |  |  |  |  |  |  |  | Y |
| “South African” | 20H/501Y.V2 |  |  |  |  |  |  |  |  |  |  |  | K |  |  |  |  | Y |
| Brazilian | 20J/501Y.V3 |  |  |  |  |  |  |  |  |  |  |  | K |  |  |  |  | Y |
| Other detected mutations |  |  |  |  |  |  |  |  | I,R,<br>G,T,<br>K | A,I,<br>R,K |  |  | Q,A,<br>D,G,<br>R,V | S,L,<br>V | L,K,<br>H,R | P,L,<br>A | H,P | R,T,<br>S |

\* The colored amino-acids are dominant (>50 %, red) or minor (<50 %, grey) at a given position. The red background highlights the emerging mutations both in clinical samples and yeast display.
\*\* The lineage designation by NextStrain initiative <sup>20</sup>; alternative strain designation proposed by Rambaut et al. <sup>21</sup> - B.1.1.7 (501Y.V1), B.1.351 (501Y.V2), and P.1 (descendent of B.1.1.28, 501Y.V3)
- Biophysical parameters of multiple selected mutant clones are shown in Table 2.

**Table 2 –.** Biophysical parameters of the mutant clones selected by yeast display.

| Clone | Library | Plasmid<sup>a</sup> | Mutations | Tm<sup>b</sup><br>[°C] | Yeast display<sup>c</sup><br>$K_{D,app}$ (pM) | Octet RED<sup>d</sup><br>$K_D$ (pM) | $k_{on}$<sup>e</sup><br>$M^{-1}s^{-1} \times 10^5$ |
| --- | --- | --- | --- | --- | --- | --- | --- |
| RBD-WT | - | pJYDC1 | AA 336 – 528 | 53.5 | $1600 \pm 200$ | $38000 \pm 10000$ | $1.7 \pm 0.06$ |
| RBD-32 | B3 | pJYDC1 | I358F, S477N, N501Y | ND | $348 \pm 35$ | ND | |
| RBD-33 | B3 | pJYDC1 | I358F, E484K, N501Y | ND | $126 \pm 2$ | ND | |
| RBD-36 | B3 | pJYDC1 | I358F, I468T, N481Y, N501Y | 54.6 | $268 \pm 3$ | ND | |
| RBD-48 | B4 | pJYDC3 | I358F, S477N, Q498R, N501Y | | $65 \pm 2$ | ND | |
| RBD-52 | B5 | pJYDC1 | I358F, N460K, E484K, S494P, Q498R, N501Y | 61.9 | $59 \pm 6$ | $3000 \pm 1000$ | $0.52 \pm 0.07$ |
| RBD-521 | B5 | pJYDC1 | I358F, N460K, E484K, Q498R, N501Y | 58.2 | $14 \pm 3$ | $340 \pm 40$ | $5.7 \pm 0.03$ |
| RBD-62 | B6(FA) | pJYDC3 | I358F, V445K, N460K, I468T, T470M, S477N, E484K, Q498R, N501Y | 57.9 | $2.5 \pm 0.2$ | $60 \pm 16$ | $13 \pm 1$ |
| RBD-71 | - | pJYDC3 | I358F, V367W, R408D, K417V, V445K, N460K, I468T, T470M, S477N, E484K, Q498R, N501Y | 63 | $8.5 \pm 1.5$ | $200 \pm 100$ | $16 \pm 1$ |

<sup>a</sup>pJYDC1 plasmid is using intrinsic eUnaG2 reporter; pJYDC3 plasmid contains DnbALFA reporter; see <sup>19</sup>
<sup>b</sup>Melting temperature as measured by differential scanning fluorimeter *Tycho NT.6* (NanoTemper Technologies GmbH)
<sup>c</sup> $K_D$ values measured between yeast surface-exposed RBD variants and the monomeric extracellular portion of ACE2 receptor.
<sup>d</sup>Measured by Octet RED96 system (ForteBio) by using AR2G biosensors. For details see Materials and Methods.
<sup>e</sup>Measured by Octet RED96 system.

Among the mutations selected and fixed in the yeast population during these initial steps of affinity maturation were three mutations that strongly emerged in clinical samples of SARS-CoV-2: S477N, E484K, and N501Y^7 8,9^. In addition, the yeast selection probed the most abundant naturally occurring variants in positions 490 and 493, which were lost in subsequent libraries of yeast selection (Table 1). S494P, which also occurs in nature but did not rapidly spread, was selected in library B4. Further analysis of this clone (Table 2, compare RBD-52 to RBD-521) shows it to increase the thermostability but decrease the association rate constant of the RBD to ACE2. The K417N/T mutations, which were identified in the “South African” and “Brazilian” variants respectively, were not included in the region mutated in libraries B3-B5, as it is distant from the binding interface.

To evaluate why some mutations were prevalent in SARS-CoV-2 and in yeast display selection, while others were not, we plotted the occurrence of all single amino-acid mutations in the GISAID database^22^ in respect to the apparent change in the RBD-ACE2 binding affinity (*K*_D,app_) as estimated by the frequency of given amino acids within a mutant library at given concentration (so-called deep mutational scanning approach^6^). Figure 2A (red and black dots) shows that the more prevalent mutations have a higher binding affinity. To quantify these results, we measured the binding of re-cloned isogenic variants of the most prevalent mutations from library B3 (Fig. 2B). The here calculated *K*_D_ values are shown as blue dots in Fig. 2A. The highest binding affinity was measured for the “South African”/”Brazilian” variant, represented by the E484K/N501Y double-mutant, which was the tightest binding clone of library B3 (Fig. 2B), followed by the “British” (N501Y) and the “European” S477N mutations (Figs. 2B and Tables 1 and 2). The *K*_D_ of the “South African” variant is 126 pM, the “British” 455 pM and for S477N a *K*_D_ of 710 pM was measured (compared to 1.6 nM for the WT). The here measured affinity data show an even stronger relation between binding affinity and occurrence in the population than can be suggested from deep-mutational scanning. To further test the lack of randomness in the selection of these mutations, we compared the occurrence of mutations for these three residues to other amino-acids in the population with their apparent binding affinity. Fig. 2C shows that indeed, in all cases, except E484R, the binding affinity of the most abundant variant in the population has the highest binding affinity at the given position. With respect to E484R, the mutation of Glu to Arg requires two nucleotide changes in the same codon, making this mutation reachable only by multiple rounds of random mutagenesis, which will delay its occurrence and may explain its low frequency. Next, we monitored whether the spread of mutation in the population also relates to the protein-stability of the RBD. Here, we used the level of yeast surface expression as a proxy to estimate the protein stability^19^. Fig. S5 shows, that mutations which occurrence is increasing in the population did not affect protein stability, corroborating this to be an important evolutionary constraint.

The most abundant naturally occurring mutations in the RBD^7–9,23^ have been selected by yeast display, already in library (B3). Next, we explored whether much higher affinity binding can be achieved by further selection.

### Exploring the affinity limits for ACE2-RBD interaction

Selection of tighter binders can demonstrate the future path of SARS-CoV-2 evolution. In parallel, an ultra-tight binder can be used as an effective ACE2 blocker, inhibiting SARS-CoV-2 infection. We used the same approach and created library B5 (Fig. 1). B5 was selected against 200, 50 and 30 pM ACE2. Sorting with less than 100 pM bait was done after overnight incubation to reach equilibrium, in 50 ml solution to prevent the ligand depletion effect (as the number of ACE2 molecules becomes much lower than the number of RBD molecules). B5 resulted in the fixation of mutations E484K, Q498R, and N501Y in all sequences clones. Mutations N460K, S477N, and S494P were present with frequencies > 20 %. Additional mutations identified were G446R, I468V, and F490Y. Representative clones with different mutational profiles were subjected to detailed analyses (Table 2).

The final selection aimed to achieve faster association-rates by using pre-equilibrium selection^24^. The new library B6(FA) (Fig. 1) was created by randomizing the RBD full-length gene, using the enriched B5 library as template. The library was pre-selected with 30 pM ACE2 for 8 hrs (reaching equilibrium after ON incubation) followed by 1 hr and 30 min incubation before selection. This resulted in the accumulation of additional mutations: V445K, I468T, T470M and also the fixation of the previously observed mutation S477N in all sequences cloned. 5 minor mutations, N354E, K417T, V367W, S494P, and S514T, with only a single sequence each, were also identified. Among them also the K417T, which is present in the “Brazilian” variant^8^. One should note that V445K and T470M require two nucleotide changes, demonstrating the efficiency of using multiple rounds of library creation on top of previous libraries (and not single clones). Interestingly, these mutations were not located at the binding interface but rather in the peripheries, which is in line with previously described computational fast association design, where periphery mutations were central^25^. From the B6(FA) library, we determined the isogenic binding for 5 different clones, with RBD-62 showing the highest affinity of 2.5 ± 0.2 pM (Fig. 2F and Tables 2 and S2). The other clones tested from the B6(FA) library have affinities between 5 to 10 pM (Fig. S3).

The ACE2 receptor and clones RBD-52, RBD-521, and RBD-62 were expressed and purified (Fig. S4). Measuring the binding affinity to ACE2 using the Octet RED96 System showed a systematically lower binding affinity in comparison to yeast titration (Table 2). For WT, yeast titration was reduced from 1.6 to 38 nM and for RBD-62 the affinity was reduced from 2.5 to 60 pM. Still, the improvement in affinity is similar for both methods (~600-fold). While most of the improvement came from reduced *k*_off_ (Fig. 2E, F) *k*_on_ increased 8-fold, from 1.7×10^5^ to 13×10^5^ M^-1^s^-1^ for RBD-62 (Table 2). In addition, RBD-62 is by 4°C more stable than the WT, probably due to the introduction of the I358F stabilizing mutation (Fig. S1). To further increase the RBD-62 affinity (stability) we prepared a site-directed mutational library on top of RBD-62, including the 15 beneficial mutations suggested from deep mutational scanning^6^, which require more than one nucleotide change to be reached (Fig. S6). None of these mutations increased the affinity towards RBD-62. Yet, a combination of three of them (V367W, R408D, and K417V) stabilized RBD-62 by 5°C, creating RBD-71, but at the cost of decreased binding affinity (Table 2, Fig. S6). This demonstrates the limitation of using single amino-acid changes from deep mutational scanning to obtain high-affinity binders.

### Structure of RBD-62 in complex with ACE2

We determined the cryo-EM structure of the N-terminal peptidase domain of ACE2 (G17-Y613) bound to RBD-62 (T333-K528, Fig. 3A and B, EMD-12187, PDB ID: 7BH9). Details of cryo-sample preparation, data acquisition, and structure determination are given in the Supplementary Materials & Methods. The cryo-EM data collection and refinement statistics are summarized in Fig. S7, S8, S9, and Table S3. Structure comparison of the ACE2-RBD-62 complex and the WT complex (PDB ID: 6M0J) revealed their overall similarity with RMSD of 0.97 Å across 586 amino acids of the ACE2 and 0.66 Å among 143 amino acids of the RBD (Fig. S10A). Three segments; R357-S371 (β2, α2), G381-V395 (α3), and F515-H534 (β11) are disordered in RBD-62, and thus not visible in the density map (blue cartoon in Fig. S10B). These segments are situated opposite to the ACE2 binding interface and therefore not stabilized and rigidified by ACE2 contacts. All mutations (except I358F) are present in the density map. Mutations V445K, N460K, I468T, T470M, S477N, E484K, Q498R, N501Y are part of the receptor-binding motif (RBM) that interacts directly with ACE2 (orange spheres Fig. 3A and B). The RBM including residues S438-Q506 shows the most pronounced conformational differences in comparison to the RBD-WT (Fig. 3C and black circle in Fig. S10A). Out of the nine mutations in the RBM, four involve intramolecular interactions, stabilizing the RBD-62 structure, including hydrogen contacts between K460 and D420, T468 and R466, and M470 and Y35. Interestingly, these are contacts gained in library B6(FA). The mutations S477N, Q498R, N501Y are forming new contacts with ACE2. Y501 makes a π interaction with Y41 and places R498 to make a hydrogen bond and salt bridge to Q42 and D38 of ACE2, forming a strong network of new interactions supporting the impact of these residues on affinity (Fig. 3D). Calculating the electrostatic potential of the RBD-62 in comparison to RBD-WT shows a much more positive surface of the former, which is complementary to the negatively charged RBD binding surface on ACE2 (Fig. 3E, F). In addition, the mutation N477 interacts with S19 of ACE2 (Fig. 3C). The interface of ACE2-RBD involves the interaction of amino acid residues from the N-terminal segment (Q24-Q42), K353, and D355 of the ACE2 domain and residues from the RBM domain of the RBD. The S477N, Q498R, and N501Y mutations in RBD-62 are situated at the two extremes of the RBD-ACE2 interface, stabilizing the complex (Fig. 3B).

**Fig. 3.**
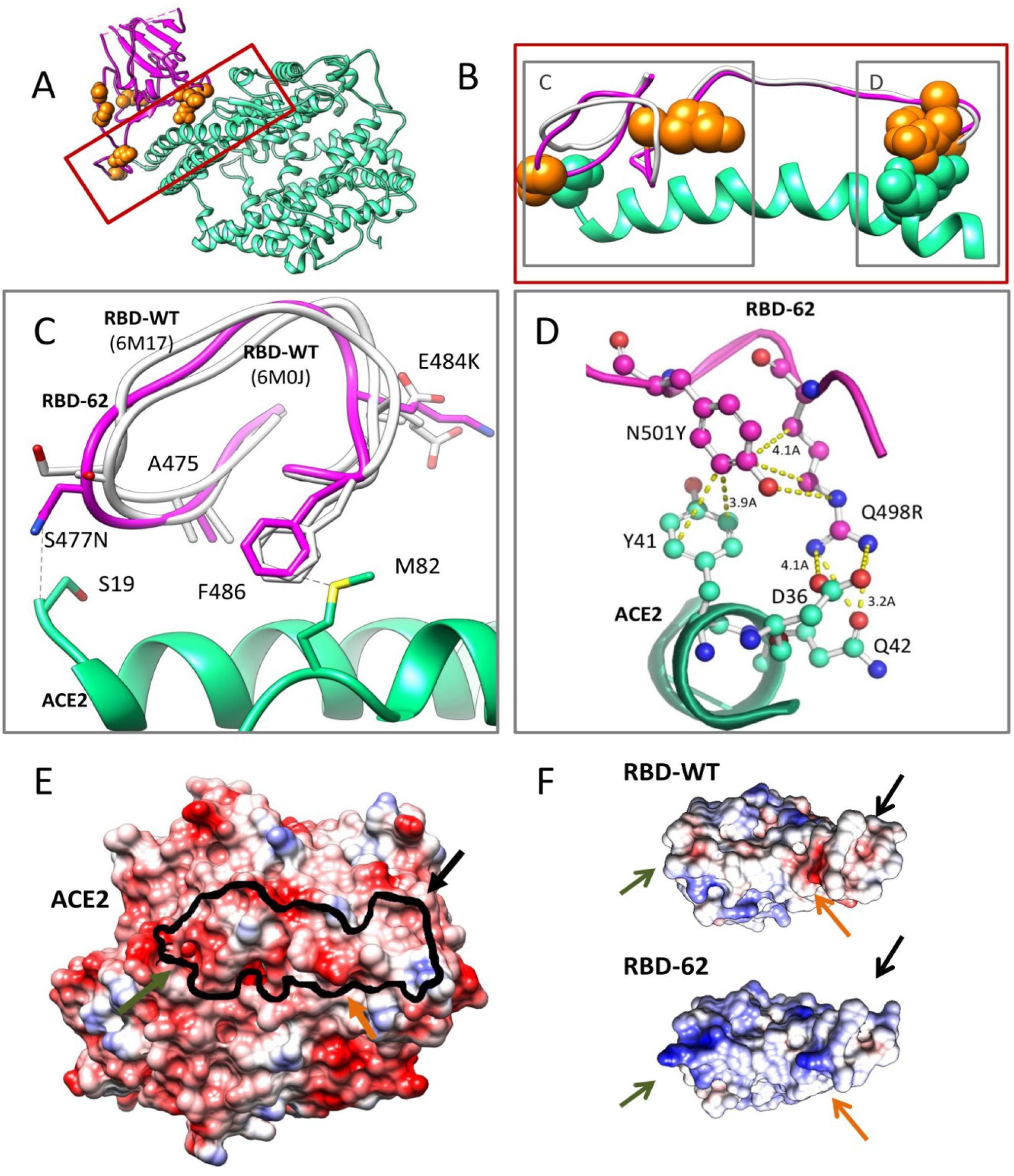
Cryo-EM structure of the ACE2-RBD-62 complex at 2.9 Å resolutions. (A) Cartoon representation of the Cryo-EM structure of ACE2 (cyan) in complex with RBD-62 RBM (magenta) with the RBD-62 mutations resolved in the density (orange spheres). (B) The S477N, Q498R, and N501Y mutations depicted in the RBM (orange spheres) interact with S19, Q42, and K353 of ACE2 respectively (cyan spheres) are situated at the two extremes of the RBD-ACE2 interface, stabilizing the complex. (C) and (D) show molecular details of selected interactions contributing to higher affinity. C) The presence of E484K and S477N causes repositioning of the RBD loop between AA475 and 487 compared to wild-type (PDB ID 6M17 and 6M0J). The change of main chain positions moves F486 to the optimum for methionine-aromatic interaction^26^. (D) The interaction network formed between RBD-62 mutations and ACE2. E) Electrostatic complementarity between RBD and ACE2 is strengthened in RBD-62 by positive charges at positions N460K, E484K, and Q498R. The black line on ACE2 indicates the RBD binding site.

### RBD-62 inhibits SARS-CoV-2 infection without affecting ACE2 enzymatic activity

A main driver of this study was to generate a tight inhibitor of ACE2 for medicinal purposes, which will be administered to the nose and lungs through inhalation. Therefore, we had to verify that the evolved RBD does not interfere with the ACE2 enzymatic activity, which is important in the Renin-Angiotensin-Aldosterone system^16,17^ We assayed the impact of RBD-WT and RBD-62 proteins on ACE2 activity. Both the *in vitro* experiment and assays done on various cells expressing ACE2 did not show differences in ACE2 activity with and without RBD-WT or RBD-62 added (Figs. 4A).

**Fig. 4.**
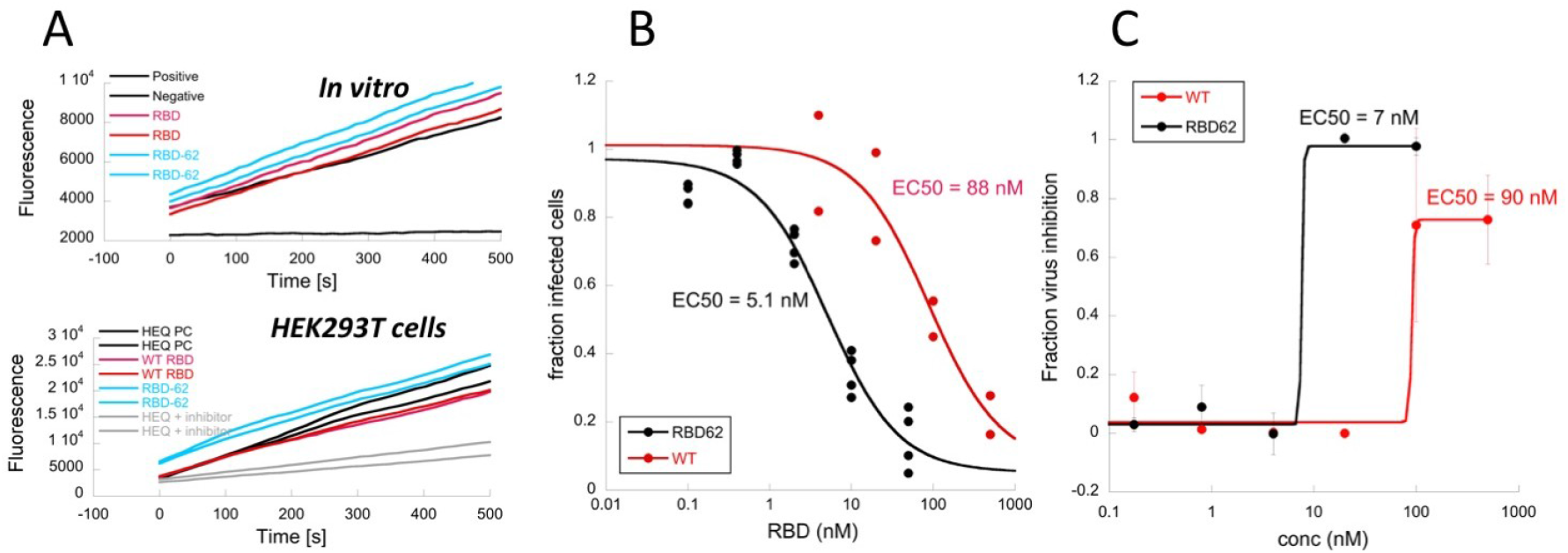
Inhibition of RBD-WT and RBD-62 on ACE2 activity and their potential to inhibit viral entry and infection. (A) ACE2 activity (*in vitro* or on cells) assayed using SensoLyte^®^ 390 ACE2 Activity Assay Kit. Fluorogenic peptide cleavage by ACE2 was measured in 10 seconds intervals over 30 minutes. The activity rate is indicated by the slope of the plot [product/time]. An ACE2 inhibitor, provided with the kit, was used as the negative control. ACE2 activity was measured *in vitro* after the addition of 100 nM of RBD-WT or RBD-62 to purified ACE2. (upper panel). ACE2 activity was measured following incubation with RBD-WT or RBD-62 on HeLa cells transiently transfected with human full-length ACE2 (bottom panel). (B) Inhibition of infection of HEK-293T cells stably expressing ACE2 by Lentivirus pseudotyped with SARS-CoV-2 spike protein. (C) Inhibition of SARS-CoV-2 infection of VeroE6 cells by RBD-WT and RBD-62 proteins.

Finally, we explored the inhibition of RBD-WT and RBD-62 on viral entry. Initially, we used Lentivirus pseudotyped with spike protein variant SΔC19^27^. Cellular entry was analyzed by flow-cytometry of Lentivirus infection promoting GFP signal. The HEK-293T cells stably expressing hACE2 were pre-incubated with serial dilutions of the two RBDs for 1 h and then the pseudovirus was added for 48 hrs. Results in Fig. 4B show that the EC50 was reduced from 88 nM for RBD-WT to 5.1 nM for RBD-62. Next, RBD-WT and RBD-62 were evaluated for their potency in inhibiting SARS-CoV-2 infection to VeroE6 cells (Fig. 4C). Similar to the pseudovirus, also here the EC50 was reduced from 90 to 6.8 nM for RBD-WT and RBD-62 respectively. More significantly, RBD-62 blocked >99% of viral entry and replication, while RBD-WT blocked only ~75% of viral replication. The complete blockage of viral replication, using a low nM concentration of RBD-62 makes it a promising drug candidate.

## Discussion

During the first year of the global SARS-CoV-2 pandemic, a small subset of mutations proved to have an adaptive advantage. First, the D614G mutation emerged and is by now prevalent. Then, the more infectious “British”, “South African”, and “Brazilian” variants appeared^28^. Among 42 non-synonymous mutations delineating the latter variants {PANGO lineages, https://www.cov-lineages.org/}, N501Y is fixed, while the K417N/T and E484K mutations are found only in the “South African” and “Brazilian” variants. Intriguingly, except for D614G, the other mutations that provide a clear advantage to the virus are located in the ACE2 binding interface of the spike protein RBD. *In vitro* evolution is a tool to directly select for higher affinity binding. Here we show that the same mutations selected by nature, S477N, E484K, and N501Y were among the first to be selected by yeast surface display affinity maturation against ACE2 (Table S4). Moreover, the E484K, N501Y variant was the tightest binding mutation emerging from the B3 library. These two mutations evolved independently, but quite rapidly merged into single clones (Table S4). The S477N mutation was selected in B3, but was fixed only in B6(FA). This is related to the smaller affinity gain of S477N in comparison to N501Y, with the double mutant E484K/N501Y increasing binding to ACE2 by the most (Table 2, S2). It is expected that *in vitro* selection follows affinity gain, but it is surprising to see that natural SARS-CoV-2 selection follows the same path. *In vitro* evolution can mimic natural evolution, on a much faster time scale. To mimic natural evolution, we pooled the selected clones from each library to construct a new one, resulting in a parallel sampling of a wider sequence-space on multiple trajectories (Fig. 1). This allowed the emergence of epistatic mutations. The use of random, rather than saturation mutagenesis is also better at mimicking natural virus evolution. RBD-62, with an affinity 600-fold higher than RBD-WT was the tightest binding clone. As seen in Table S4, many mutations were transients along the selection process, while a small number persisted (as is observed also for SARS-CoV-2). Q498R was first observed in library B4, but established itself only in B5. This mutation emerges only together with N501Y, suggesting an epistatic effect between the two mutations. Indeed, deep-mutational scanning of single RBD mutations^6^ showed Q498R to negatively affect both protein stability and binding, while we show that in combination with N501Y binding affinity is improved by ~4-fold above N501Y alone. The epistatic effect is also confirmed by the RBD-62 ACE2 complex structure, where Y501 is placing R498 to form multiple interactions. Single mutation scanning does not account for this kind of cooperativity, which has a crucial role within protein-protein interfaces^29^. The rapid spread of N501Y in the population increases the likelihood for the emergence of the Q498R mutant, which will probably have even higher infectivity. Conversely to Q498R, both the N501Y and E484K mutations established themselves independently, allowing for their rapid *in vitro* selection and emergence in SARS-CoV-2. This suggests that with the spread of the “British”, “Brazilian”, and “South African” variants, we project that the Q498R mutation will appear in the future, on top of these mutations. The synergism of Q498R with N501Y and E484K increases ACE2 binding by ~50-fold relative to WT. Another mutation projected to follow is N460K, which emerged late in yeast display. These mutations are located in a hyper-variable region of the RBD (Fig. S11), suggesting that their appearance is not constrained.

Infectivity is one concern of the emerging mutations, but equally important is their potential for immune evasion, both from the resistance provided by previous infections of the WT virus, and even more importantly, from vaccination. To evaluate the effect of these RBD mutations on antibody binding, we manually inspected 92 antibody-RBD (nanobody, Spike) structures for clashes, replacing the WT RBD with the new RBD-62 structure, as determined in complex with ACE2 using Cryo-EM. 28 of the antibodies bind outside the RBM and 8 interactions are not affected by the mutations on RBD-62. However, for 56 antibodies the interaction was compromised and for 9 antibodies, major clashes with RBD-62 were identified (Fig. S12). Notably, E484K and Q498R caused most of the observed effects. While E484K is now prevalent, Q498R has not yet been identified in patients.

An intriguing question is whether the spreading of the tighter binding SARS-CoV-2 variants in humans is accidental. From the similarity to yeast display selection, where stringent conditions are used, one may hypothesize that stringent selection is also driving the rapid spread of these mutations. Abundant low-quality face masks may provide one such selection condition, as they reduce viral titer, but not sufficiently. Therefore, higher quality face-masks (N95) should be encouraged, particularly in closed environments.

The RBD-62 has the potential as a drug, as it blocks ACE2 with very high affinity, while preserving the ACE2 functionality. It completely inhibited SARS-CoV-2 infection of VeroE6 cells, at low nM concentration. Further experiments are now on the way to evaluate the efficacy of RBD-62 as a drug. The main advantage of blocking ACE2 is that it is not directly affected by viral escape mutations, which constantly evolve. Despite the high hopes that vaccines will eradicate COVID-19, realistically, the development of a working drug stays high on the agenda.

## Conclusions

Knowing the enemy provides an advantage to the defender. Using *in vitro* evolution, we successfully delineated the pathway of SARS-CoV-2 towards higher infectivity. While SARS-CoV-2 may evolve using other criteria (such as shown for the D614G mutation of the spike protein), we show that currently, the strongest evolutionary driver towards higher infectivity is the increase in RBD-ACE2 binding affinity. This is in line with the higher infectivity of SARS-CoV-2 relative to SARS-CoV. Therefore, it is important to sequence the *RBD* of SARS-CoV-2 for a larger number of patients, which will provide early identification of upcoming more infective variants. Moreover, knowing the future evolution of the virus allows time to test the efficacy of vaccines and drugs against those variants already now. Meanwhile, N95 masks should be promoted as the new variants are more infectious. Finally, the methods applied in this study could be generalized for other viruses, once their evolutionary path is identified.

## Acknowledgments

We thank Dr. Shira Albeck, Dr. Tamar Unger, and Dr. Yoav Peleg from the structural proteomics unit (ISPC) of the Weizmann Institute of Science for their invaluable help in cloning and protein production. We thank Dr. Jaroslav Nunvar and CB2 group for valuable discussion and support.

## Authors contribution

J.Z. and G.S. conceived the project; J.Z., S.M., M.S., E.Z., and G.S. performed experiments; N.E. prepared cryo-EM samples and built atomic models and refined structures with O.D. J.Z, N.E, O.D, J.R and G.S wrote the manuscript.

## Data and materials availability

Cryo-EM data have been deposited in the Electron Microscopy Data Bank with accession codes EMD-12187, PDB ID: 7BH9.

## Supplementary Materials

## Materials and methods

### Cloning and DNA manipulations

The RBD domain variants (see Table S1) were PCR amplified (KAPA HiFi HotStart ReadyMix, Roche, Switzerland) from codon-optimized SARS-CoV-2 Spike protein gene (Sino Biological, SARS-CoV-2 (2019-nCoV) Cat: VG40589-UT, GenBank: QHD43416.1) by using appropriate primers. Amplicons were purified by using NucleoSpin^®^ Gel and PCR Clean-up kit (Nacherey-Nagel, Germany) and eluted in DDW. Yeast surface display plasmid pJYDC1 (Adgene ID: 162458) and pJYDC3 (162460) were cleaved by *Nde*I and *Bam*HI (NEB, USA) restriction enzymes, purified, and tested for non-cleaved plasmids *via* transformation to *E. coli* Cloni^®^ 10G cells (Lucigen, USA). Each amplicon was mixed with cleaved plasmid in the ratio: 4 μg insert: 1 μg plasmid per construct, electroporated in *S. cerevisiae* EBY100^30^, and selected by growth on SD-W plates.

Cloning of ACE2 extracellular domain (AA G17-Y613) gene and *RBDs* into vectors pHL-sec^31^ were done in two steps. Initially, the RBD gene was inserted in helper vector pCA by restriction-free cloning^32^. pCA is a pHL-sec derivative lacking 862 bp in the GC rich region (nt 672 – 1534). In the second step, the correctly inserted, verified by sequencing, *RBDs* with flanking sequences were cleaved by using restriction enzymes *Xba*I and *Xho*I (NEB, USA) and ligated (T4 DNA ligase, NEB, USA) in cleaved full-length plasmid pHL-sec.

Site-directed mutagenesis of *RBDs* was performed by restriction-free cloning procedure^32^. Megaprimers were amplified by KAPA HiFi HotStart ReadyMix (Roche, Switzerland), purified with NucleoSpin™ Gel and PCR Clean-up Kit (Nachery-Nagel, Germany), and subsequently inserted by PCR in the destination using high fidelity Phusion^®^ (NEB, USA) or KAPA polymerases. The parental plasmid molecules were inactivated by *Dpn*I treatment (1 h, NEB, USA) and the crude reaction mixture was transformed to electrocompetent *E. coli* Cloni^®^ 10G cells (Lucigen, USA). The clones were screened by colony PCR and their correctness was verified by sequencing.

### DNA libraries preparation

SARS-CoV-2 RBD gene (RBD) libraries were prepared by MnCl_2_ error-prone mutagenesis^33^ using Taq Ready-mix (Hylabs, Israel). The mutagenic PCR reactions (50 μl) were supplemented with increasing MnCl_2_ concentrations: 0.05, 0.1, 0.2, 0.4, 0.6, 0.8 and 1.0 nM. Template DNA concentration ranged between 100 and 400 ng per reaction and 20 – 30 reaction cycles were applied. The amplified DNA was purified, pooled, and used directly for yeast transformation via electroporation. The whole gene randomization amplicon comprised the whole RBD gene (AA 336-528, pJYDC1 vector). Libraries B3, B4, and B5 were prepared by homologous recombination of an invariant fragment of RBD with necessary overlaps (AA 336-431) and the mutagenized library fragment (AA 431-528). The mutagenic fragments were prepared by the same error-prone PCR procedure (20 cycles).

### Yeast transformation, cultivation, and expression procedures

The detailed description of all the procedures and our enhanced yeast display platform itself was described in details^19^. Briefly, plasmids were transformed into the EBY100 *Saccharomyces cerevisiae*^30 34^. Single colonies were inoculated into 1.0 ml liquid SD-CAA media and grown overnight at 30°C (220 rpm). The overnight cultures were spun down (3000 g, 3 min) and the exhausted culture media was removed before dilution in the expression media 1/9^19^ to OD ~ 1. The expression cultures were grown at different temperatures 20, 30, and 37 °C for 8 – 24 h at 220 rpm, depending on the experimental setup. The expression co-cultivation labeling was achieved by the addition of 1 nM DMSO solubilized bilirubin (pJYDC1, eUnaG2 reporter holo-form formation, green/yellow fluorescence (Ex. 498 nm, Em. 527 nm)) or 5 nM ALFA-tagged mNeonGreen (pJYDC3, DnbALFA). Aliquots of cells (100 ul) were collected by centrifugation (3000 g, 3 min) resuspended in ice-cold PBSB buffer (PBS with 1 g/L BSA), passed through cell strainer nylon membrane (40 μM, SPL Life Sciences, Korea), and analyzed.

### Binding assays and affinity determination using yeast surface display

Aliquots of yeast expressed and labeled cells ready for flow-cytometry analysis were resuspended in analysis solution with a series of labeled ACE2 concentrations. The concentration range was of CF®640R succinimidyl ester labeled (Biotium, USA) ACE2 extracellular domain (AA Q18 – S740) was dependent on the protein analyzed (0.1 pM – 50 nM). The analysis solution volume was adjusted (1 – 100 ml) to avoid the ligand depletion greater than 10% as well as the time needed to reach the equilibrium (1 h – 12 h, 5 rpm, 4 °C)^35^. After the incubation, samples were collected (3000 g, 3 min), resuspended in 200 ul of ice-cold PBSB buffer (200 μl), passed through a cell strainer, and analyzed. The expression and binding signals were determined by flow cytometry using BD Accuri™ C6 Flow Cytometer (BD Biosciences, USA). The cell analysis and sorting were done by S3e Cell Sorter (BioRad, USA). The analysis was done by single-cell event gating (Fig. S2), green fluorescence channel (FL1-A) was used to detect RBD expression positive cells (RBD+) via eUnaG2 or DnbALFA, and far-red fluorescent channel (FL4-A) recorded CF®640R labeled ACE2 binding signals (CF640+). The eUnaG2 signals were automatically compensated by the ProSort™ Software and pJYDNp positive control plasmid (Adgene ID 162451^19^). The mean FL4-A fluorescence signal values of RBD+ cells, subtracted by RBD-, were used for the determination of binding constant *K*_D_. The standard non-cooperative Hill equation was fitted by nonlinear least-squares regression using Python 3.7. The total concentration of yeast exposed protein was fitted together with two additional parameters describing the given titration curve^6^.

### Production and purification of RBD and ACE2 proteins

The extracellular part of ACE2 (Q18 – S740) and RBD protein variants (Table S1) were produced in Expi293F cells (ThermoFisher). Pure DNA was transfected using ExpiFectamine 293 Transfection Kit (ThermoFisher) using the manufacturer protocol. 72 hours post-transfection, the cells were centrifuged at 1500 rpm for 15 minutes. The supernatant was filtered using 0.45 μm Nalgene (ThermoFisher) filter and the pellet was discarded. The filtered supernatant was loaded onto a 5 ml of HisTrap Fast Flow column (Cytivia (GE, USA), cat 17-5255-01). ÄKTA pure (Cytivia, USA) was used to purify the protein. The column was washed in 25 mM Tris, 200 mM NaCl 20 mM imidazole, then, the protein was eluted using gradient elution with elution buffer containing 25 mM Tris, 200 mM NaCl 1M imidazole. Buffer exchange to PBS and the concentration of the protein were done by using amicons® (Merck Millipore Ltd, cat:UFC900324).

### Cryo-Electron Microscopy

#### Sample preparation

2.5 μl of ACE2-RBD-62 complex at 3.5 mg/ml concentration was transferred to glow discharged UltrAuFoil R 1.2/1.3 300 mesh grids (Quantifoil), blotted for 2.5 seconds at 4°C, 100% humidity, and plunge frozen in liquid ethane cooled by liquid nitrogen using a Vitrobot plunger (Thermo Fisher Scientific).

#### Cryo-EM image acquisition

Cryo-EM data were collected on a Titan Krios G3i transmission electron microscope (Thermo Fisher Scientific) operated at 300 kV. Movies were recorded on a K3 direct detector (Gatan) installed behind a BioQuantum energy filter (Gatan), using a slit of 20 eV. Movies were recorded in counting mode at a nominal magnification of 165,000x, corresponding to a physical pixel size of 0.53 Å. The dose rate was set to 16.2 e^-^/pixel/sec, and the total exposure time was 1.214 sec, resulting in an accumulated dose of 70 e^-^/Å^2^. Each movie was split into 57 frames of 0.021 sec. The nominal defocus range was −0.7 to −1.1 μm, however, the actual defocus range was larger. Imaging was done using an automated low dose procedure implemented in SerialEM^36^. A single image was collected from the center of each hole using image shift to navigate within hole arrays and stage shift to move between arrays. The ‘Multiple Record Setup’ together with the ‘Multiple Hole Combiner’ dialogs were used to map hole arrays of up to 3×3 holes. Beam tilt was adjusted to achieve coma-free alignment when applying image shift.

#### Cryo-EM image processing

Image processing was performed using CryoSPARC software v3.0.1^37^. The processing scheme is outlined in Fig. S7. A total of 4470 acquired movies were subjected to patch motion correction, followed by patch CTF estimation. Of these, 3357 micrographs having CTF fit resolution better than 5 Å and relative ice thickness lower than 1.07, were selected for further processing. Initial particle picking was done using the ‘Blob Picker’ job on a subset of 100 micrographs. Extracted particles were iteratively classified in 2D and their class averages were used as templates for automated particle picking from all selected micrographs, resulting in 2,419,995 picked particles. Particles were extracted, binned 6×6 (60-pixel box size, 3.18 Å/pixel), and cleaned by multiple rounds of 2D classification, resulting in 1,649,355 particles. These particles were used for *ab initio* 3D reconstruction with 5 classes. Out of the 5 classes only one, containing 552,575 particles, refined to high resolution. Two additional classes may show ACE2 in a closed conformation (containing 249,841 and 503,670 particles), however, they did not refine, partially because of preferred orientation. The 3D class containing 552,575 particles was refined as follows: Particles were re-extracted only from micrographs with defocus lower than 1.7 μm, binned 2×2, and subjected to homogeneous refinement (355,891 particles, 200-pixel box size, 1.06 Å/pixel). The particles were then sub-classified into 2 classes, and particles from the higher-resolution class were re-extracted without binning in 680-pixel boxes, subjected to per particle motion correction, followed by non-uniform refinement^38^ with per-particle defocus optimization. The final map, at a resolution of 2.9 Å (Fig. S8), was sharpened with a B-factor of −83 before atomic model building. In the final map, the RBD is only partially resolved at the distal region from the ACE2 interface. To better understand the reason for the missing density, we subjected the particles from the well-refined 3D class (355,891 particles) to variability analysis^39^, with a binary mask imposed on the RBD region (Fig. S9). Classification into 5 distinct classes based on 3 eigenvectors, revealed variable density at the RBD distal region, which could not be modeled reliably. The cryo-EM data collection process and refinement statistics are summarized and visualized in Fig. S7, S8, S9, and Table S3.

#### Model building

The atomic model of the ACE2-RBD-62 was solved by docking into the CryoEM maps the homologous refined structure of the SARS-CoV-2 spike receptor-binding domain bound with ACE2 (PDB-ID 6M0J) as a model, using the Dock-in-Map program in PHENIX^40^. All steps of atomic refinements were carried out with the Real-space refinement in PHENIX^41^. The model was built into the cryo-EM map by using the COOT program^42^. The ACE2-RBD-62 model was evaluated with the MOLPROBIDITY program^43^. The ACE2 (G17-Y613) contains one zinc ion linked to H374, H378, and E402 and three N-acetyl-β-glucosaminide (NAG) glycans linked to N53, N90, and N546. In the RBD-62 structure (T333-K528) three fragments; R357-S371 (β2, α2), G381-V395 (α3), and F515-H534 (β11) are disordered, and thus not visible in the density map. Details of the refinement statistics of the ACE2-RBD62 structure are described in Table S3. 3D visualization and analyses were performed using UCSF Chimera^44^ and PyMol (Schrödinger, Inc.; 2.4.0).

### Analysis of RBD circulating virus variants

All amino acid substitutions in the RBD (116) were downloaded from the GISAID database (23 December 2020)^22^ with the corresponding numbers of sequences and regions and plotted against the binding (ΔLog_10_(K_D, App_)) or expression (ΔLog_10_MFI) extracted from the RBD deep mutational scanning dataset^6^. We gratefully acknowledge all GISAID contributors and Starr et all for sharing their data.

### Octet RED binding analysis

Octet RED96 System (forte BIO, Pall Corp., USA) was used for real-time binding determination. Briefly, 10 μg/ml of ACE2 diluted in 10 mM NaAcetate pH5.5 was immobilization to an amine-reactive 2G biosensor using standard procedure. The purified RBD was diluted in a sample buffer (PBS+0.1% BSA+0.02% Tween20). Analyte concentrations, association, and dissociation times were adjusted per sample. Data Analysis v10 software (forte BIO, Pall Corp., USA) was used for data fitting, with the mathematical model assuming a simple 1:1 stoichiometry.

### Pseudo-virus production and inhibition of infection by RBD

Pseudo-virus production: SARS-CoV-2-Spike pseudotyped Lentivirus was produced by cotransfection of Hek293T cells pCMV ΔR8.2, pGIPZ-GFP, (*26*) and pCMV3 SΔC19 at a ratio of 1:1:1. 24 hours before the transfection 1 x 10^6^ cells were seeded into a 10 cm plate. On the day of the transfection cells were washed by Dulbecco’s Modified Eagle’s Medium (DMEM) (Gibco 11965092) and 5 ml of Opti-MEM (Gibco 11058021) was added to the plate. 10 μg of plasmids mix was transfect using lipofectamine 2000 transfection reagent (Thermo Fisher 11668027) according to the manufacturer’s instructions. After 4 hours, the media was replaced by 9 ml of fresh media. The supernatant was harvested 72 h post-transfection, centrifuged (1000 g, 5 min), and filtered to remove all residual debris (Millex-HV Syringe Filter Unit, 0.45 μm).

RBD inhibition assay: HEK-293T cells stably expressing hACE2 (GenScript M00770) were seeded into a 24-well plate at an initial density of 6 x 10^4^ cells per well. The following day cells were pre-incubated with serial dilutions of RBDs (1 h) and then the pseudotyped Lentivirus was added. After 24 h, the cell medium was replaced with fresh DMEM, and cells were grown for an additional 24 h. After this procedure, cells were harvested and the GFP signal was analyzed by flow cytometry (BD Accuri™ C6 Plus Flow Cytometer, BD Biosciences, USA).

### Inhibition of SARS-CoV-2 infection

The strain 2019-nCoV/IDF0372/2020 was supplied by the National Reference Centre for Respiratory Viruses hosted by Institute Pasteur (Paris, France) and headed by Dr. Sylvie van der Werf. The human sample from which strain 2019-nCoV/IDF0372/2020 was isolated has been provided by Dr. X. Lescure and Pr. Y. Yazdanpanah from the Bichat Hospital. The experiments were done by Institute Pasteur. VeroE6 (C1008) cells were grown in DMEM with 10% serum and 1% penicillin to 50% confluence in 384-well format and incubated with RBDs at given concentration for 2 hrs before 0.1 MOI of SARS-CoV-2 was added for one hour. The inoculum was subsequently removed and a medium with the RBD was added. After 48 hrs of incubation, the supernatant was recovered and viral load was measured using RT-PCR with a forward primer: TAATCAGACAAGGAACTGATTA, reverse primer: CGAAGGTGTGACTTCCATG. In parallel, cell viability was assessed after 48 hrs incubation using the CellTiter Glo kit from Promega. Raw data are normalized against appropriate negative and positive controls and are expressed as the fraction of virus inhibition. The curve fit was performed using the variable Hill slope model of four parameters logistic curve:

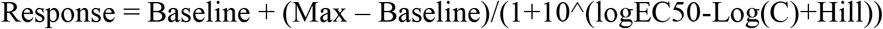

### ACE2 activity assay

Human ACE2 activity was evaluated using SensoLyte® 390 ACE2 Activity Assay Kit (ANASPEC; cat# 72086) according to manufacturer’s protocol, with the following changes – assay was performed in 384 well plates with a ratio of 1:5 of the recommended volume of buffer, substrate, and inhibitor. The activity was measured on either purified ACE2 (0.75 ng; Abcam, aB351852) or the following cell lines – HeLa transiently transfected with ACE2 (6000 cells per assay), HEK-293T stable transfected with ACE2 (GenScript M00770, 8000 cells per assay), Caco2 cells (40,000 cells per assay). 10 nM WT RBD and RBD-B62 were added before activity measurement. The activity rate is indicated by the slope of the plot [product/time].

### Defining the RBD domain for yeast display and protein expression

To optimize the RBD for yeast display affinity maturation, we screened multiple different constructs for yeast surface expression. RBDs of different starting and termination positions were cloned in a pJYDC1 vector and their impact on expression, stability, and ACE2 binding were determined (Table S1). The RBDcon1 was the shortest construct lacking the last C-terminal loop of the RBD domain (516 – 528) and including one unpaired cysteine. This resulted in poor expression and binding. The RBDcon2 and con3 included this loop, resulting in domain stabilization and an increase both in binding and expression. Although RBDcon4 (published by Kramer et al^45^) construct demonstrated high expression yields both in yeast and Expi293F™ cells, as well as good thermo-stability, we decided not to use it in yeast display since one unpaired cysteine (C538) is close to its C-terminus and the construct contains part of the neighboring domain. We continued with the RBDcon2 and RBDcon3 constructs for yeast display and protein expression in Expi293F™ cells respectively.

**Table S1 –.** Comparison of different RBD domain spans and their influence on domain properties

| Construct | Position<sup>a</sup> | Number of AA | Size [kDa] | Yeast expression [mean FL1*10<sup>3</sup>]<sup>b</sup> | Yeast display estimated $K_D$ [nM]<sup>c</sup> | Melting temperature [°C] |
| --- | --- | --- | --- | --- | --- | --- |
| RBDcon1 | 336-516 | 181 | 20.5 | 16.4 | 3.2 ± 0.1 |  |
| RBDcon2 | 336-528 | 193 | 21.7 | 32.9 | 1.6 ± 0.3 |  |
| RBDcon3 | 333-528 | 196 | 22.1 | 37.9 | 1.1 ± 0.3 | 53.5 ± 0.3 |
| RBDcon4 | 319-541 | 223 | 25.1 | 67.6 | 1.2 ± 0.2 | 53.8 ± 0.2 |

<sup>a</sup> numbers are according to UniProtKB- P0DTC2
<sup>b</sup> measured in pJYDC1 (eUnaG2 fluorescence signal)
<sup>c</sup> Binding affinity against ACE2 was determined by FACS, with the relevant construct expressed on yeast surface.

**Table S2 –.**
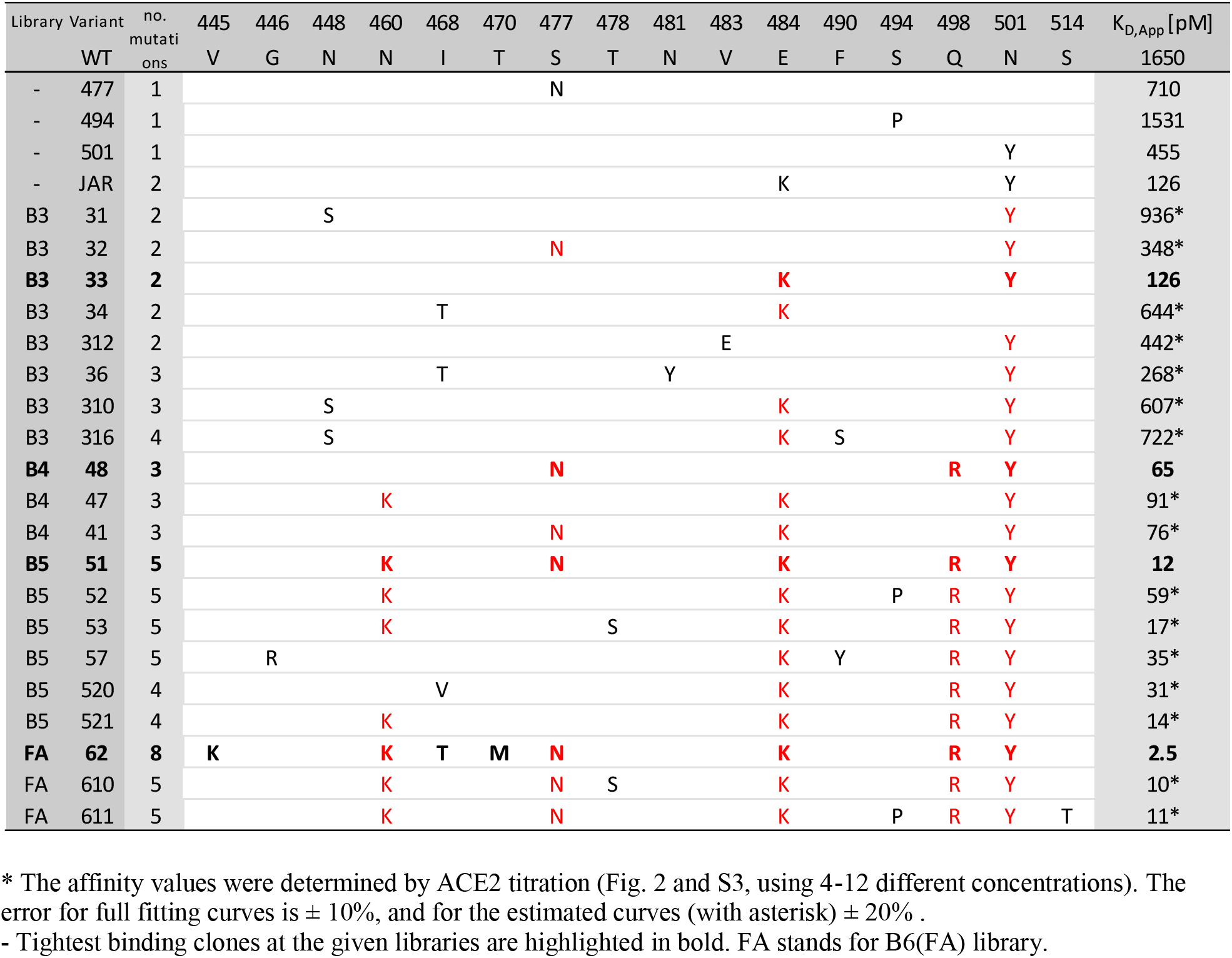
RBM of analyzed mutant clones selected by yeast display.

**Table S3 –.**
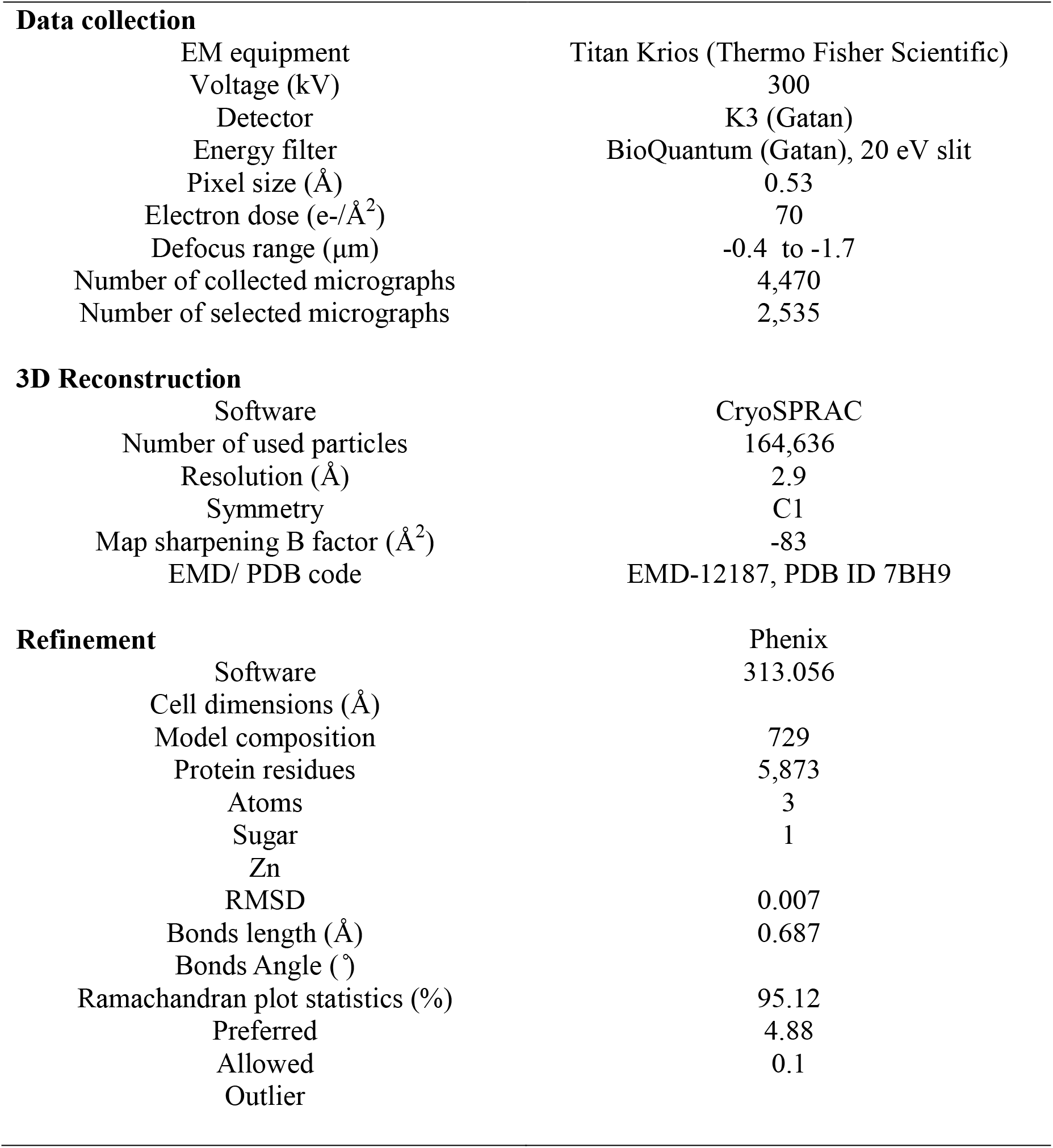
Cryo-EM data collection and refinement statistics of ACE2-RBD62 complex

**Table S4 –.**
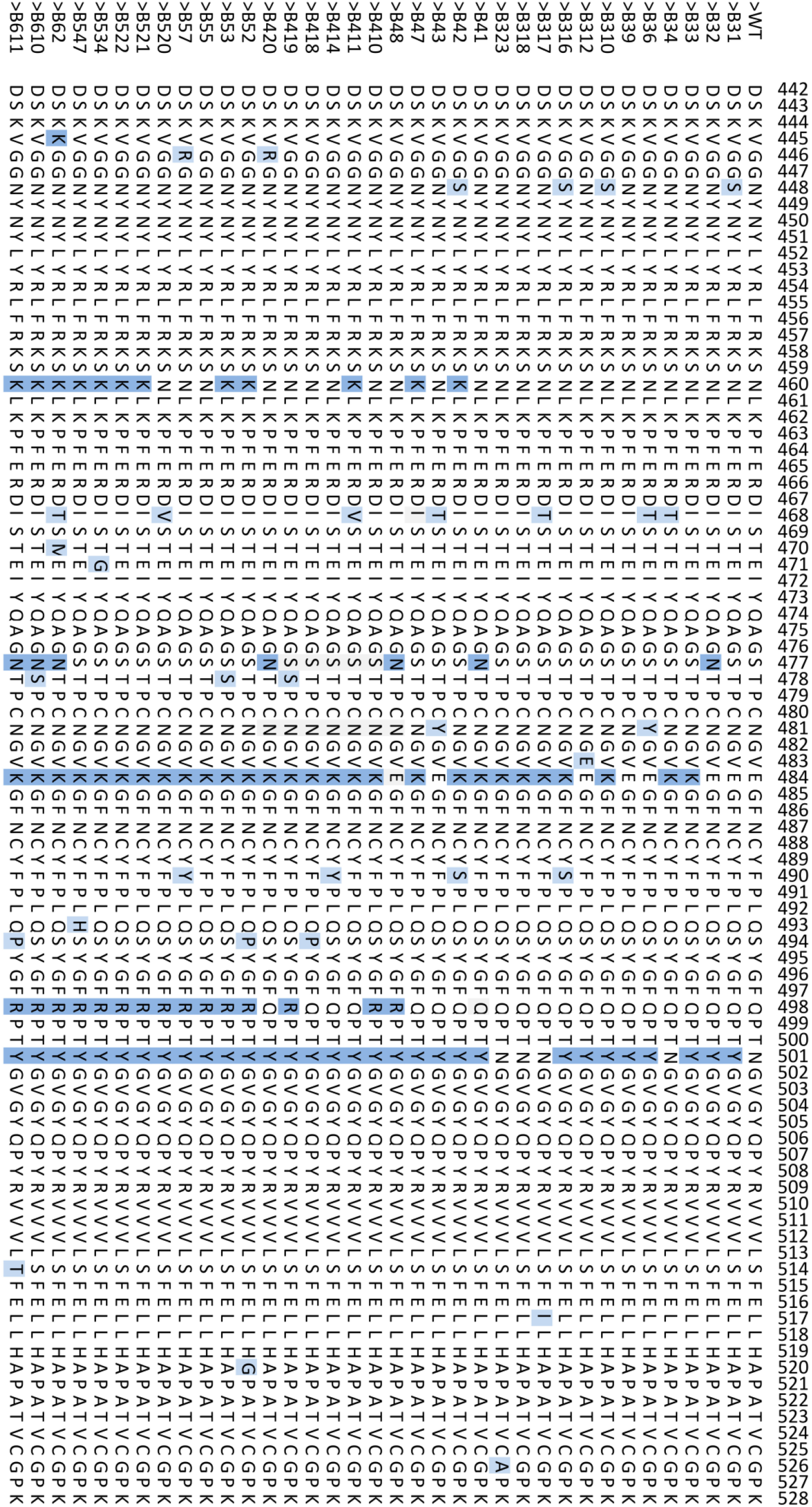
Non-redundant clones sequenced during the RBD affinity maturation process. Rare/transient mutations are in light blue. Important changes are highlighted in dark blue.

### Supplementary Material Figures

**Fig. S1.**
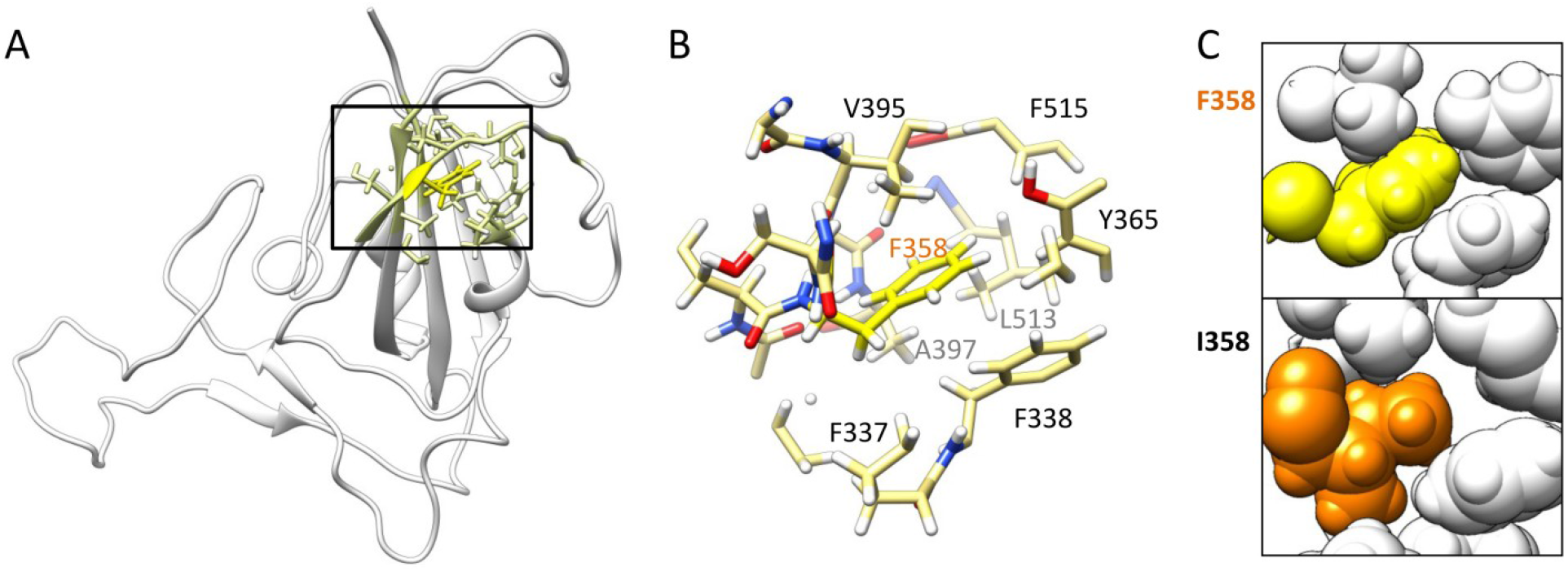
The I358F mutation, selected by yeast surface display, increases protein stability and expression. (A) The position of I358F (bright yellow) mutation in the RBD structure (PDB ID 6M17) and the neighboring residues within 5 Å distance (pale yellow). (B) Shows the residues involved in the formation of the hydrophobic cavity around the I358F mutation predicted from the X-ray structure. Additional residues that are involved: K356, R357, S359, V395, Y396. (C) The spherical representation of predicted phenylalanine mutant position inside the hydrophobic cavity (yellow) and wild-type residue (isoleucine, orange).

**Fig. S2.**
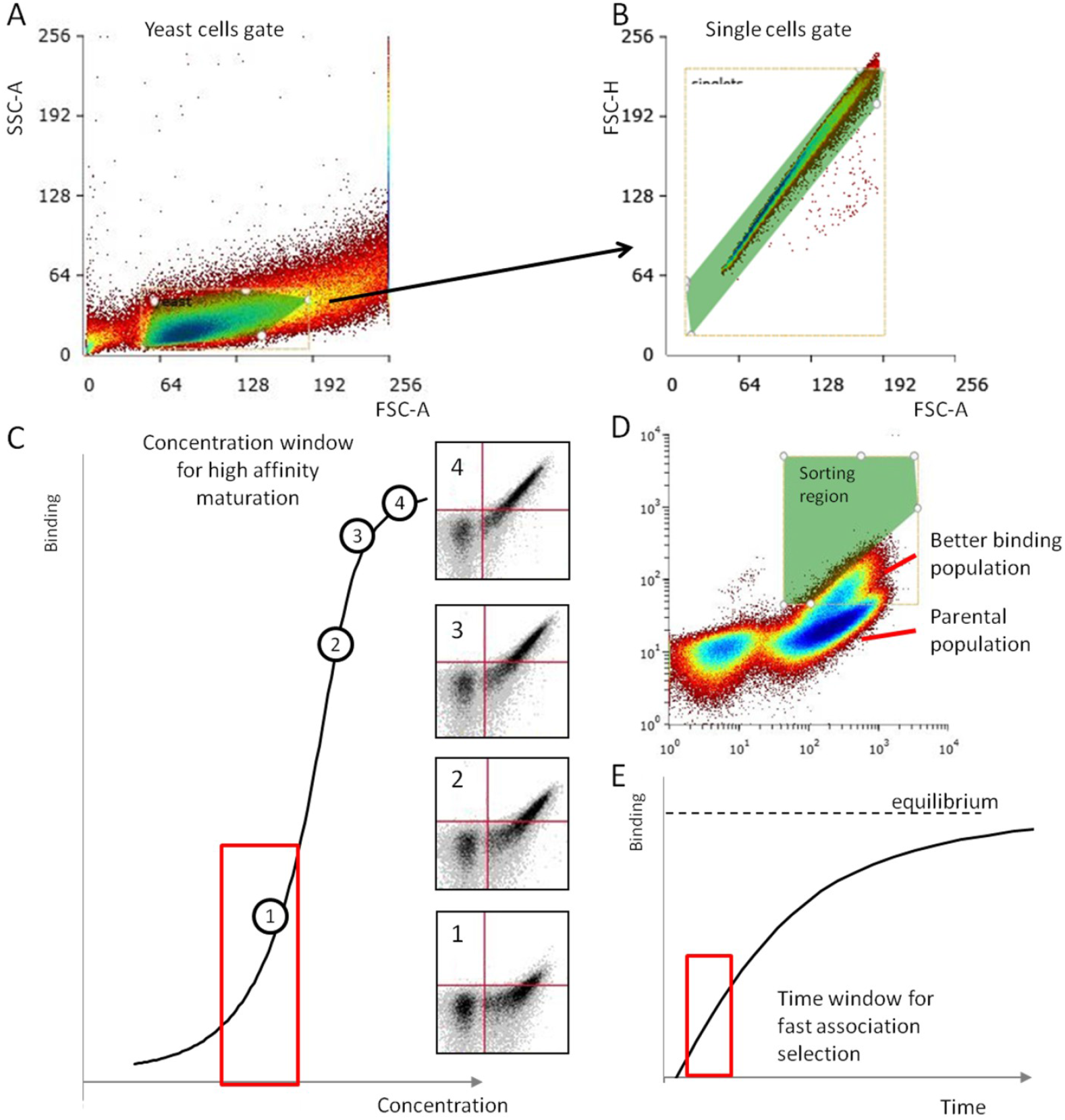
Gaiting and selection strategies for *in vitro* evolution of SARS-CoV-2 RBD domain. (A, B) Gating strategy for FACS sorting. In the first step, yeast cells were isolated by their FSC-A and SSC-A properties (A). In the second step (B), single cells were isolated by their FSC properties (area and height) on the diagonal plot. The Green area represents the gated region. C) Selection strategy for affinity maturation. The library was titrated with a range of ACE2 concentrations to select the concentration with limited signal (inset 1). Under such conditions, the tighter binding clones gain the highest advantage over the parental population. Using less stringent selection (insets 2 – 4) reduces the advantage of the tighter binders. Using too low concentrations of ACE2 protein will also result in loss of selectivity. (D) Affinity maturation library after 3 sorts, where the separation between parental and tighter binding population is well defined. The top 0.1 – 0.3 % of cells were sorted – green region. (E) Fast association selection strategy. The library was incubated with a constant concentration (30 pM) of ACE2 for different times. The time with minimal signal was determined and used for the selection of clones with faster association (library B6(FA)). The same shape of the sorting region as in (D) was applied.

**Fig. S3.**
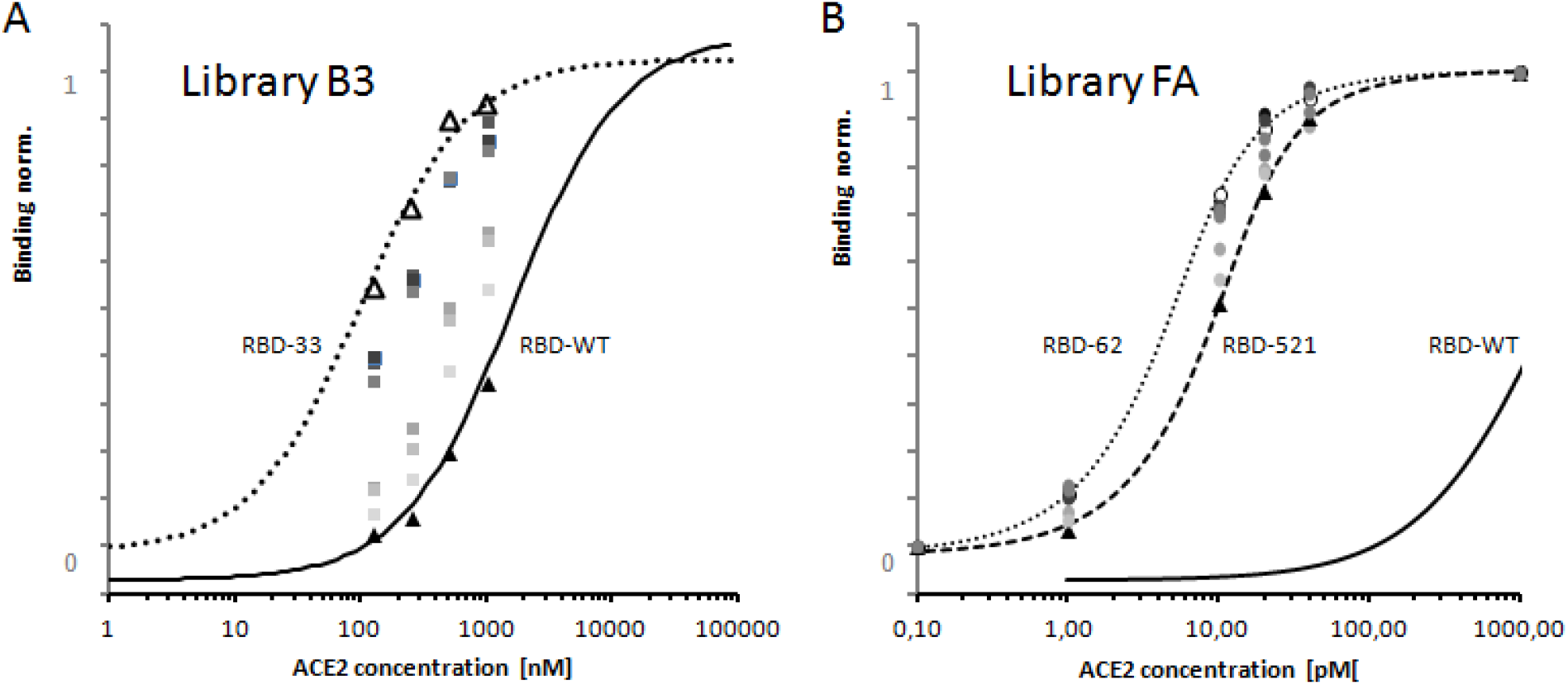
Evaluating the binding affinity of selected clones from libraries B3 and B6(FA). Five single-clones were evaluated for binding to ACE2 from each library, to determine the range of affinity maturations after FACS selection. Each clone was incubated with four (library B3) or six (B6FA) different concentrations of ACE2. The binding curve was fitted using two additional parameters describing the curve (range, baseline).

**Fig. S4.**
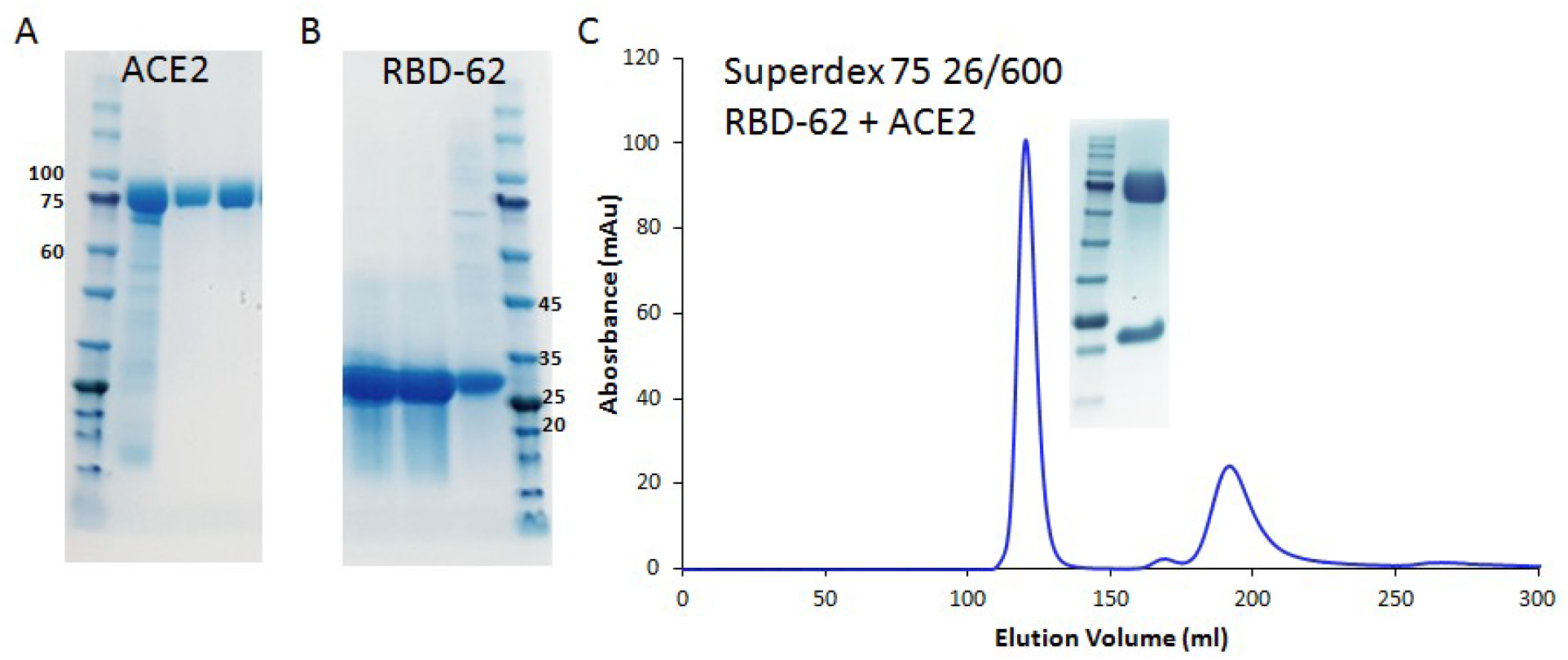
Protein purification of ACE2, RBD-62 and the complex between the two. Both proteins were expressed in Expi293F cells and secreted to cell culture media. (A) SDS-PAGE analysis after NiNTA agarose purification of ACE2 receptor extracellular domain (AA Q18 – S740). (B) SDS-PAGE analysis from NiNTA agarose purification of RBD-62 (AA 333-528). (C) The ACE2 + RBD-62 complex was purified by gel filtration chromatography column prior to CryoEM. ACE2 protein was mixed with an excess of RBD-62 (1:1.5), incubated 1h on ice, and applied on the chromatography column by using ÄKTA pure FPLC system. The first peak corresponds to the complex (SDS-gel inset) and the second peak represents excess RBD-62.

**Fig. S5.**
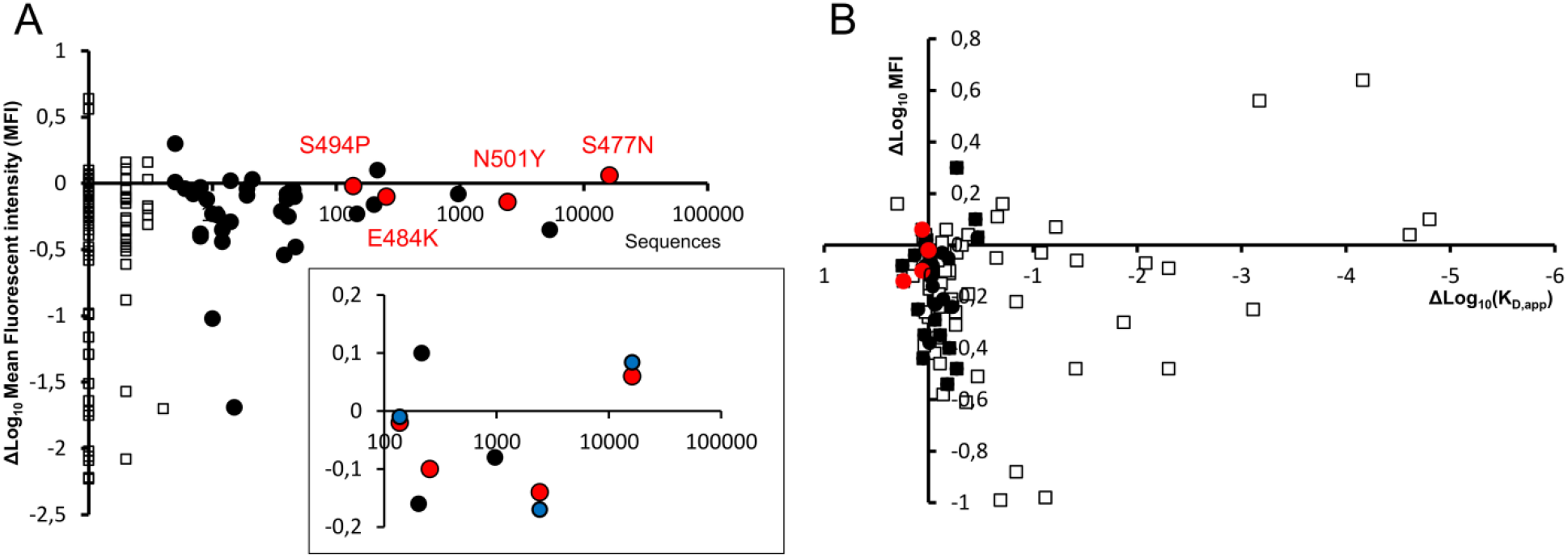
SARS-CoV-2 RBD mutations frequencies in the population and their level protein expression. (A) Relation between the impact of mutations on yeast surface expression and their occurrence in the population. Expression was measured as the mean fluorescence intensity (MFI) of the specific clone expressed on the yeast surface by Star ret al.^6^ (black and red) or by us (blue, inset). Empty squares and black dots are showing data with < 5 or > 5 sequences recorded, respectively. The emerging mutations in the population are shown in red. The graph shows that the variance in expression decreases with higher occurrence in the population. (B) Relation between the affinity (x-axis), expression (y-axis), and the occurrence in population: Empty squares < 5 sequences; black dots > 5 sequences; red dots represent four emerging mutations (all with more than 100 sequences). Based on (A) and (B), rapidly spreading mutations are affinity enhancing without compromising the protein stability.

**Fig. S6.**
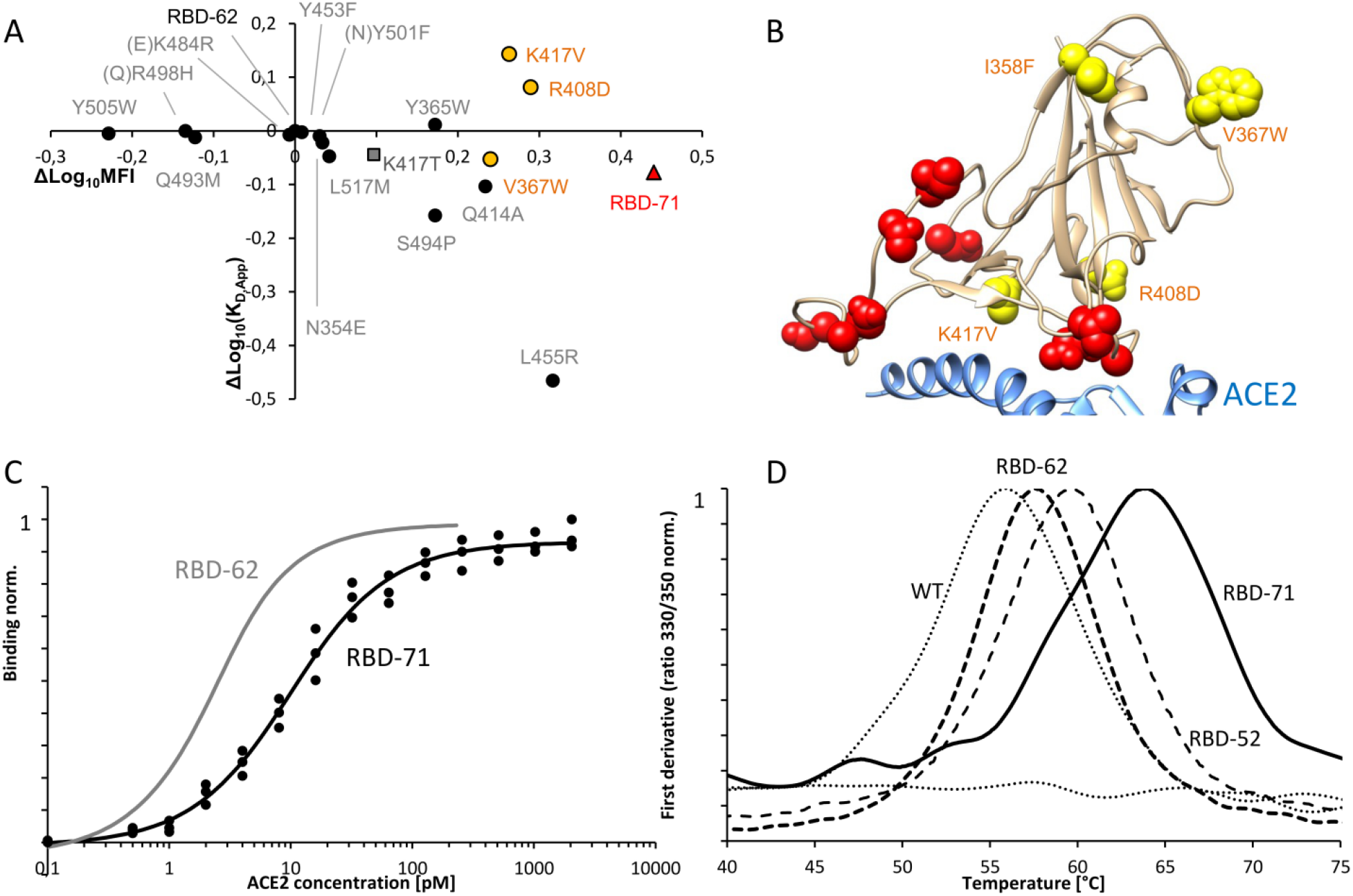
Site-directed mutagenesis of RBD-62, using the affinity enhancing mutations. 15 mutations were predicted to enhance RBD-ACE2 binding or stability^6^. Mutation K417T (grey) was introduced from the B4(FA) library. These mutations were evaluated for enhancing the affinity of RBD-62 towards ACE2 and their effect on stability. (A) Impact of mutations, on top of RBD-62 on ACE2 binding (y-axis) and yeast surface expression. Three mutations (orange circles), which have the highest impact on expression, were combined in RBD-71 (red triangle). (B) Localization of stabilizing (yellow) and binding enhancing mutations depicted in the RBD structure (PDB ID 6m17, best rotamer is shown). (C) Binding curve of RBD-71 with RBD-62 for comparison. (D) Normalized protein melting curves for RBD-WT, RBD-62, RBD-52, and RBD-71 measured using the Tycho NT.6 (NanoTemper).

**Fig. S7.**
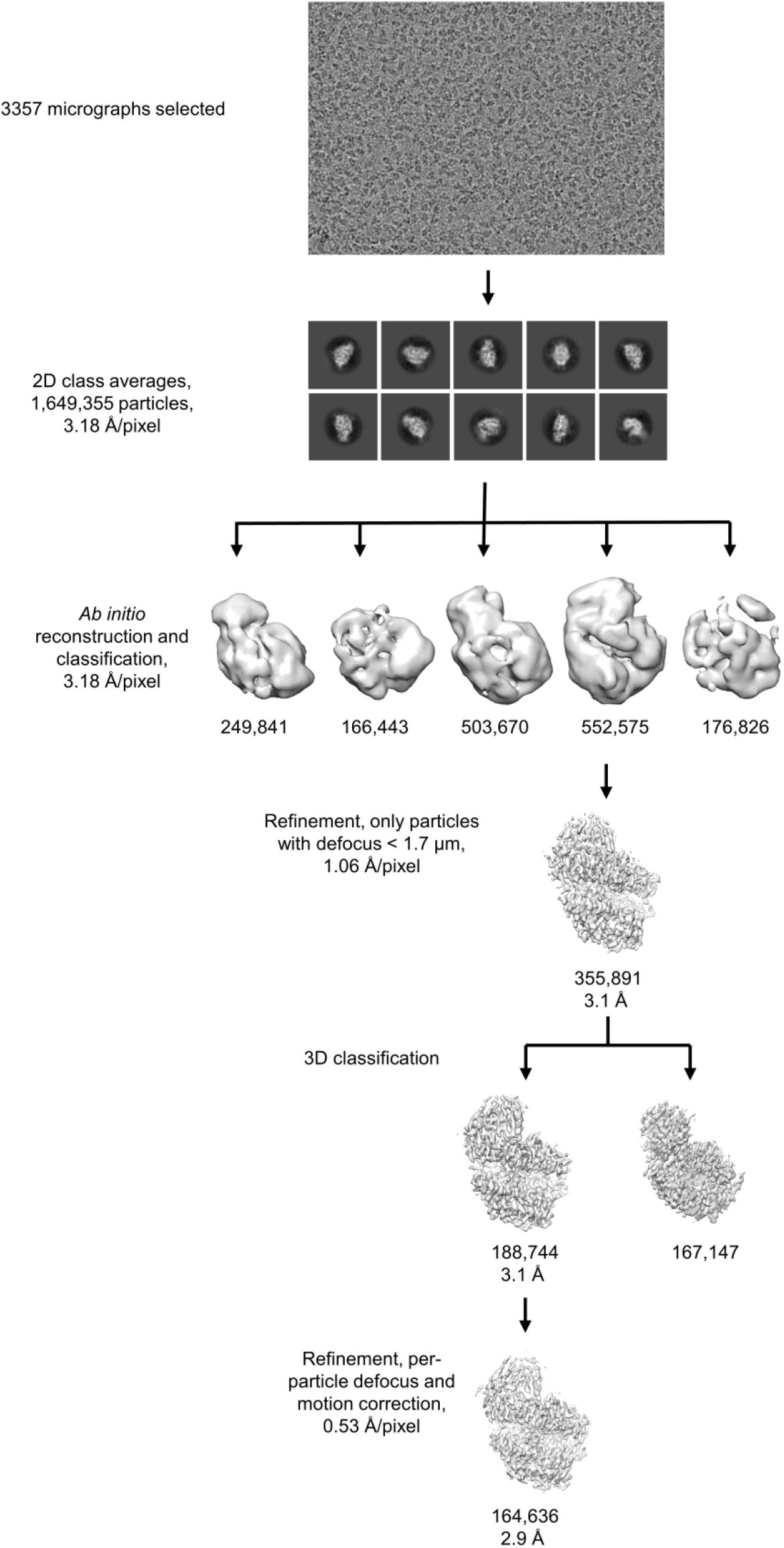
Single-particle cryo-EM processing scheme. The details of the process are described in the Methods section under “Cryo-EM image processing”. The number of particles in each map is indicated under the map’s image, along with the map’s resolution where relevant.

**Fig. S8.**
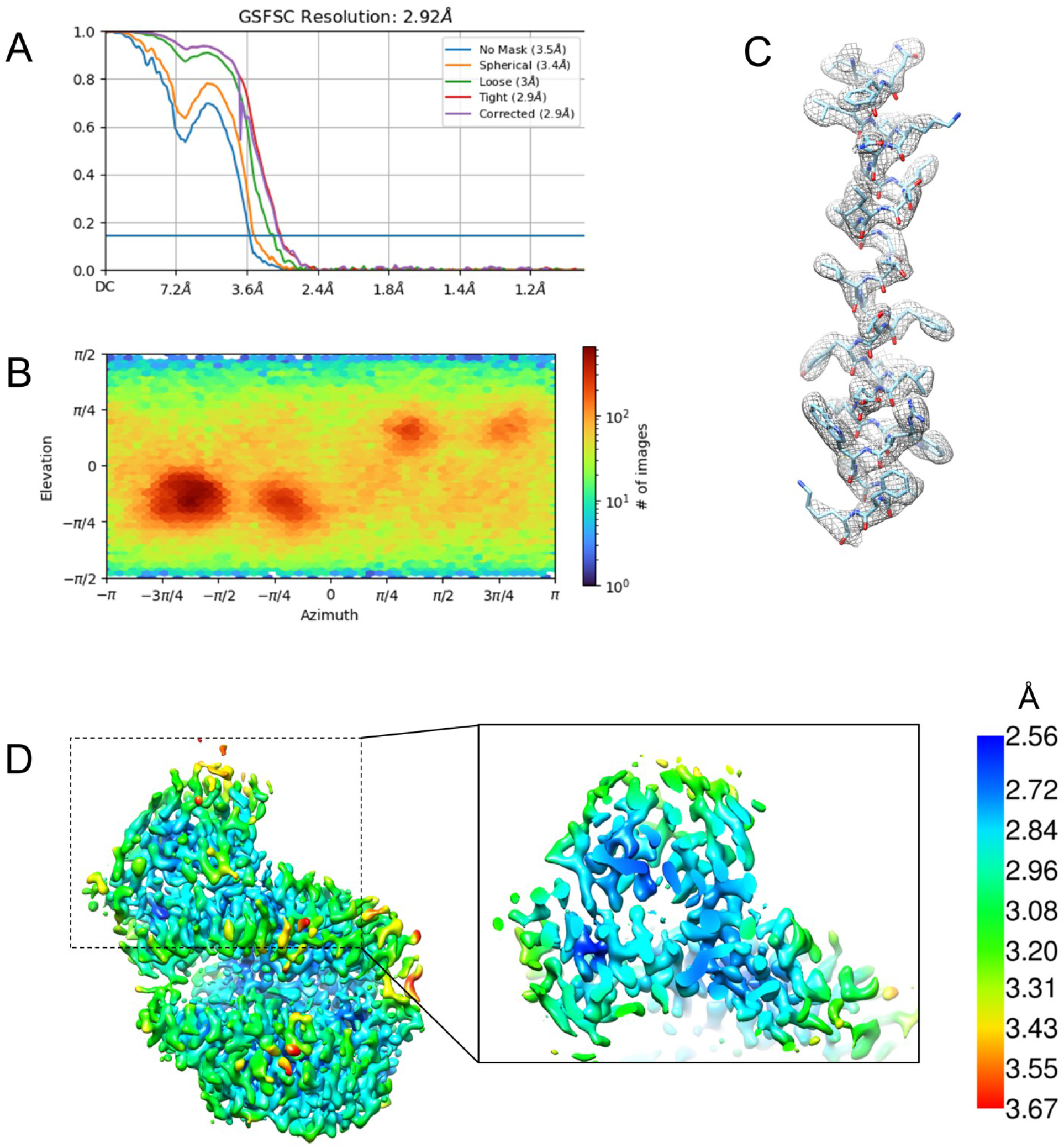
Resolution estimate and angular distribution for the ACE2-RBD-62 cryo-EM map. (A) Fourier Shell Correlation (FSC) curves. (B) Angular distribution plot. (C) An alpha-helical segment showing the map density and fitted atomic coordinates. (D) Cryo-EM map colored according to local resolution estimate. The inset shows a slice through the RBD-ACE2 interface.

**Fig. S9.**
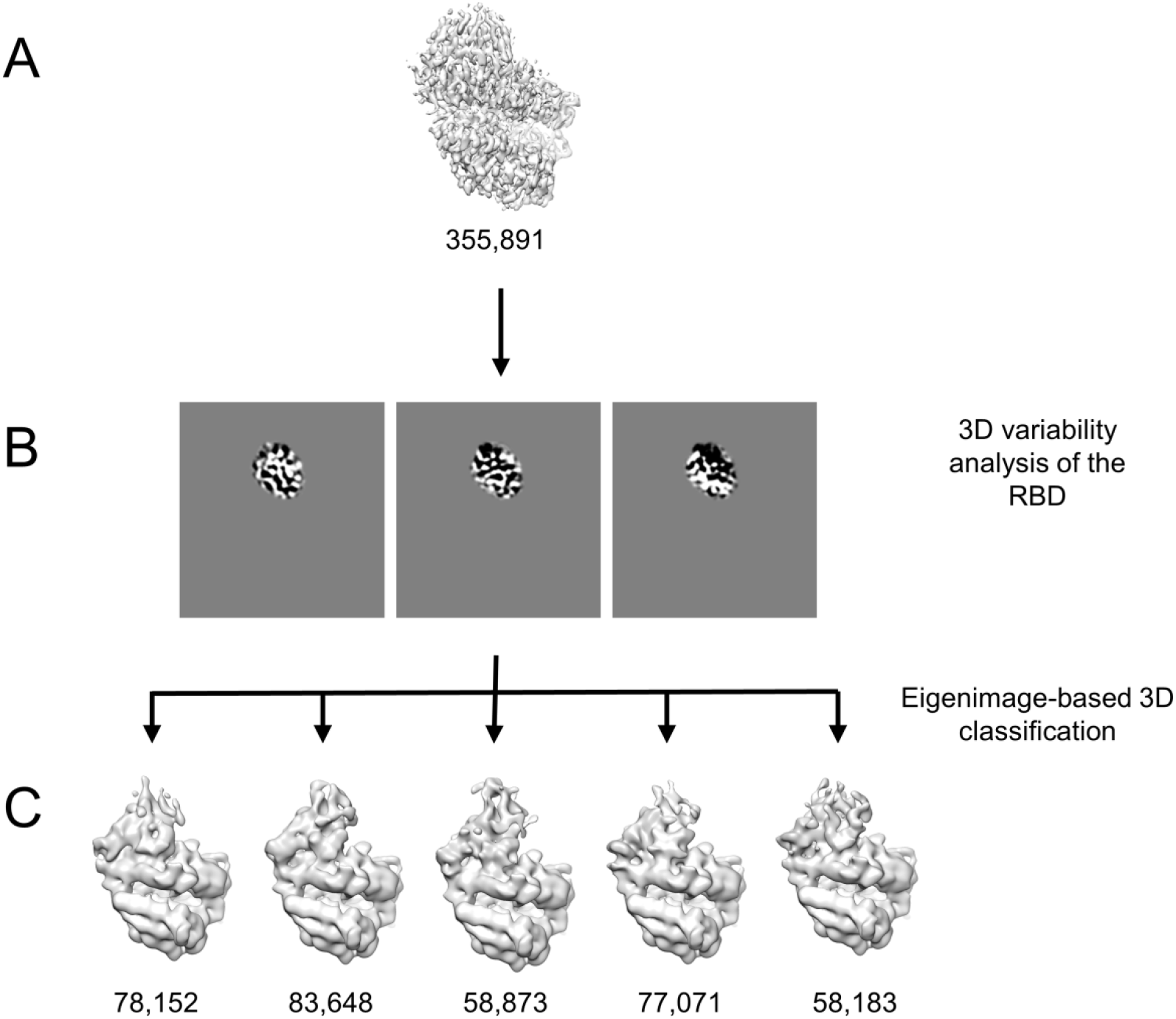
Variability analysis of the RBD. (A) Particle images from the well-resolved 3D class were subjected to 3D variability analysis. (B) Central slices through the three eigenimages calculated with a binary mask around the RBD region. (C) Five 3D classes, which were calculated based on the eigenimages. The maps show variable density for the RBD.

**Fig. S10.**
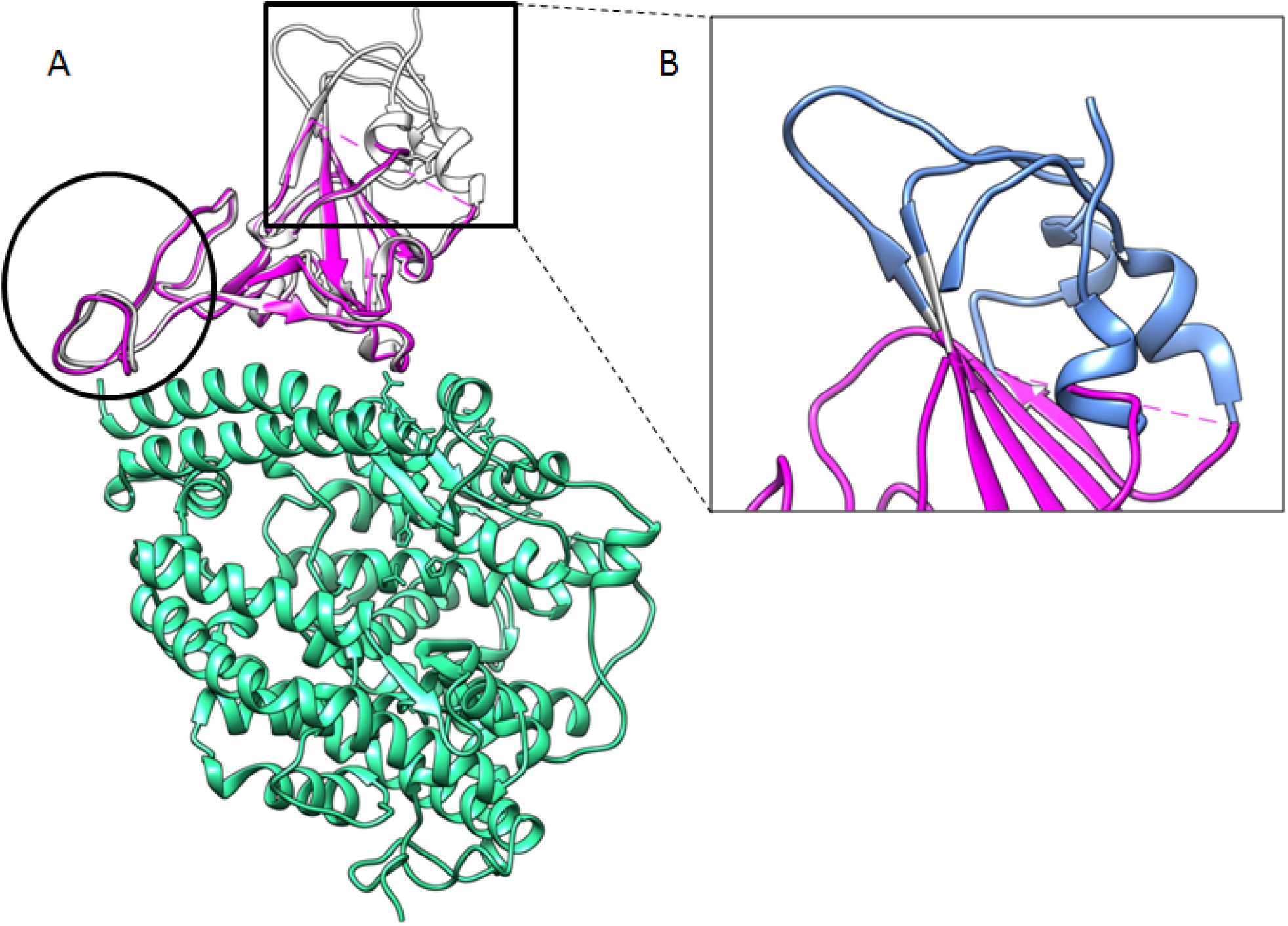
Global comparison between RBD-WT and RBD-62 shows overall similarity. (A) RBD-62 preserves its typical twisted five-stranded antiparallel β sheet (β1, β3-β5, and β10) with an extended insertion containing the short β5-β9 strands, α4, and η3 helices and loops. The biggest differences are pronounced between M470 and F490 (black circle). (B) The upper part comprising of three segments: R357-S371 (β2, α2), G381-V395 (α3), and F515-H534 (β11) is not resolved in the density map (blue ribbon, added from PDB ID: 6M0J).

**Fig. S11.**
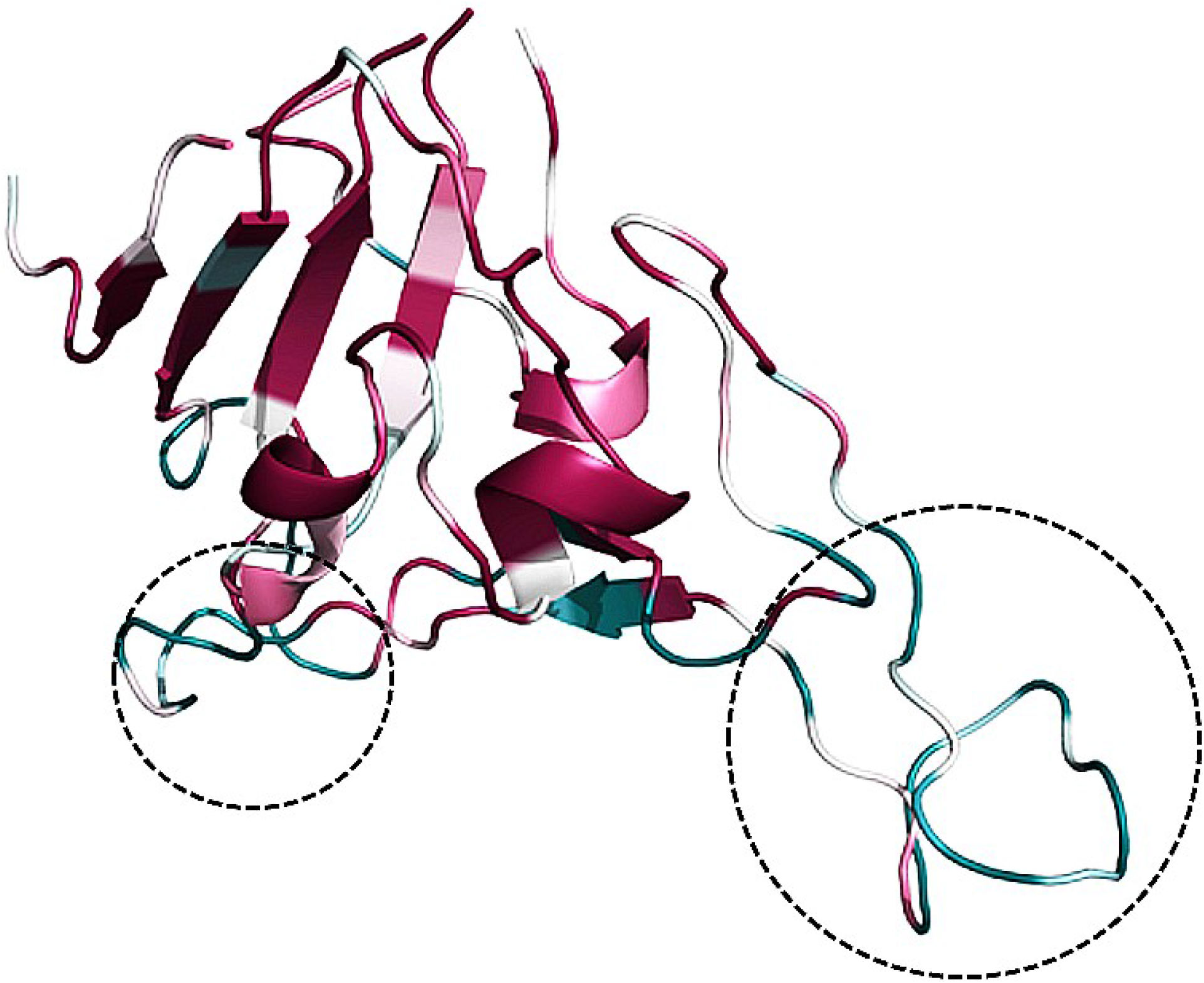
An analysis of conserved positions computed by ConSurf server depicted on the RBD-62 structure. The amino acids are colored by their conservation grades with turquoise-through-maroon indicating variable-through-conserved by ConSurf server^46^.

**Fig. S12.**
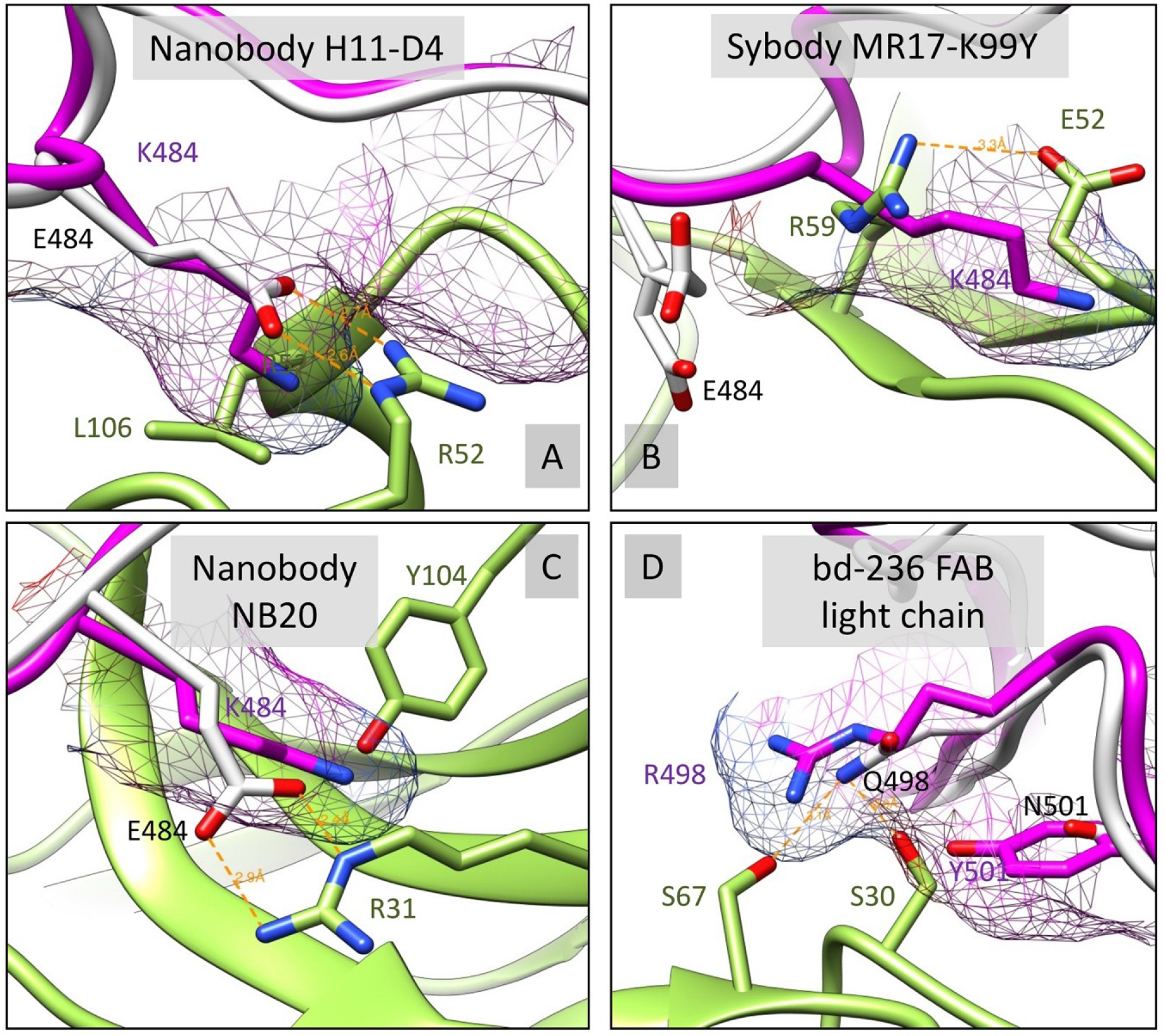
RBD-62 mutations are interfering with binding to multiple antibodies. The RBD-62 (magenta, mesh surface) was structurally overlaid with RBD-WT (white). S477N, E484K, Q498R, and N501Y RBD mutated residues were analyzed for disruption of wild-type contacts (orange dashed line) and clashes with corresponding binding antibody/nanobody (green) in relation to RBD-WT. Four examples A) PDB ID: 6YZ5, B) PDB ID: 7CAN, C) PDB ID: 7JVB, D) PDB ID: 7CHE were the mutations resulted in a dramatic impact are shown.

